# A neural model of proximity to reward

**DOI:** 10.1101/2022.10.03.510669

**Authors:** P. Botros, N. Vendrell-Llopis, R. M. Costa, J. M. Carmena

## Abstract

Throughout learning, refinement of cortical activity in cortex, a process termed “credit assignment”, underlies the refinement of behavioral actions leading to reward. While previous research shows striatum’s role in linking behavior to reward, striatum’s role in linking the underlying behaviorally-relevant cortical activity to reward remains unclear. Leveraging a neuroprosthetic task while recording from the rat cortex and striatum, we demonstrate that the striatum encodes the dynamics of the proximity of cortical activity to reward. Such encoding was independent from external task feedback and emerged as cortical activity consolidated over learning, with dorsal and ventral striatum playing complementary yet distinct roles. Striatal activity thus constitutes a neural model of cortical progress towards reward, suggesting one mechanism by which the brain implements credit assignment to refine behavior.

## Main

When we first begin learning a new skill, initial behavior is highly variable as we explore which actions lead to reward (*1*). As behavioral variability decreases with training, neural activity patterns in cortex that relate to those behaviors simultaneously reduce in their variability (*2, 3*). The computational theory underlying this process of neural consolidation during skill learning is frequently termed the “credit assignment problem”: to learn, the brain must discover and subsequently bias task-relevant cortical neurons to execute intended behavior more consistently (*4*), despite reinforcement signals being both spatially non-specific (*5*) and temporally delayed (*6, 7*). While particular subcortical areas have been identified as necessary or sufficient for credit assignment (*8–10*), a full understanding of the neural basis of skill learning remains elusive.

As animals attempt to maximize reward over time throughout learning, a solution to the credit assignment problem inevitably involves the relationship between ongoing behavior, relevant cortical activity, and proximity to reward. To this end, myriad studies suggest the striatum as a strong candidate for a home for this relationship: the striatum is necessary for skill learning (*9, 11, 12*), encodes reward-related quantities (*13*), and connects cortex and reward-relevant subcortical areas (*14*). Existing studies suggest different roles during learning across the dorsal/ventral axis of the striatum (*11, 15*) and across cell types (*16*), predicting heterogeneity in striatal encoding. However, it remains unclear how striatum might facilitate credit assignment to task-relevant cortical neurons and how such a role might differ between striatal regions.

Taking advantage of a neuroprosthetic learning task (*8–10*) in which observed neural activity causally determines behavior, we studied how the striatum models the relationship between task-relevant cortical activity and reward and how this encoding emerges over learning. We simultaneously recorded tens to hundreds of single units from the motor cortex, dorsomedial striatum (DS), and ventral striatum (VS) of 7 rats over 8-15 days of training (n=80 days total, Figure 1A-B, Supplementary Figure 1) using chronically implanted Neuropixels 1.0 electrodes (*17*). Each day, we divided four well-isolated units located in the motor cortex, henceforth designated as “direct units”, into two ensembles, *E1* and *E2*, and defined a simple decoder that computed the difference between the summed firing rates within each ensemble (“E1 - E2”, Figure 1C). The rat’s goal was to modulate cortical activity such that the decoder output exceeded either a positive (T1) or negative (T2) target (Figure 1D-F). This “hit” yielded either smaller or larger amounts of sucrose water reward, depending on target (Figure 1D-F). All cortical units other than direct units were designated as “indirect units”. During trials, an auditory tone with its frequency derived from the decoder provided external feedback to rats of task progress.

**Figure 1.**
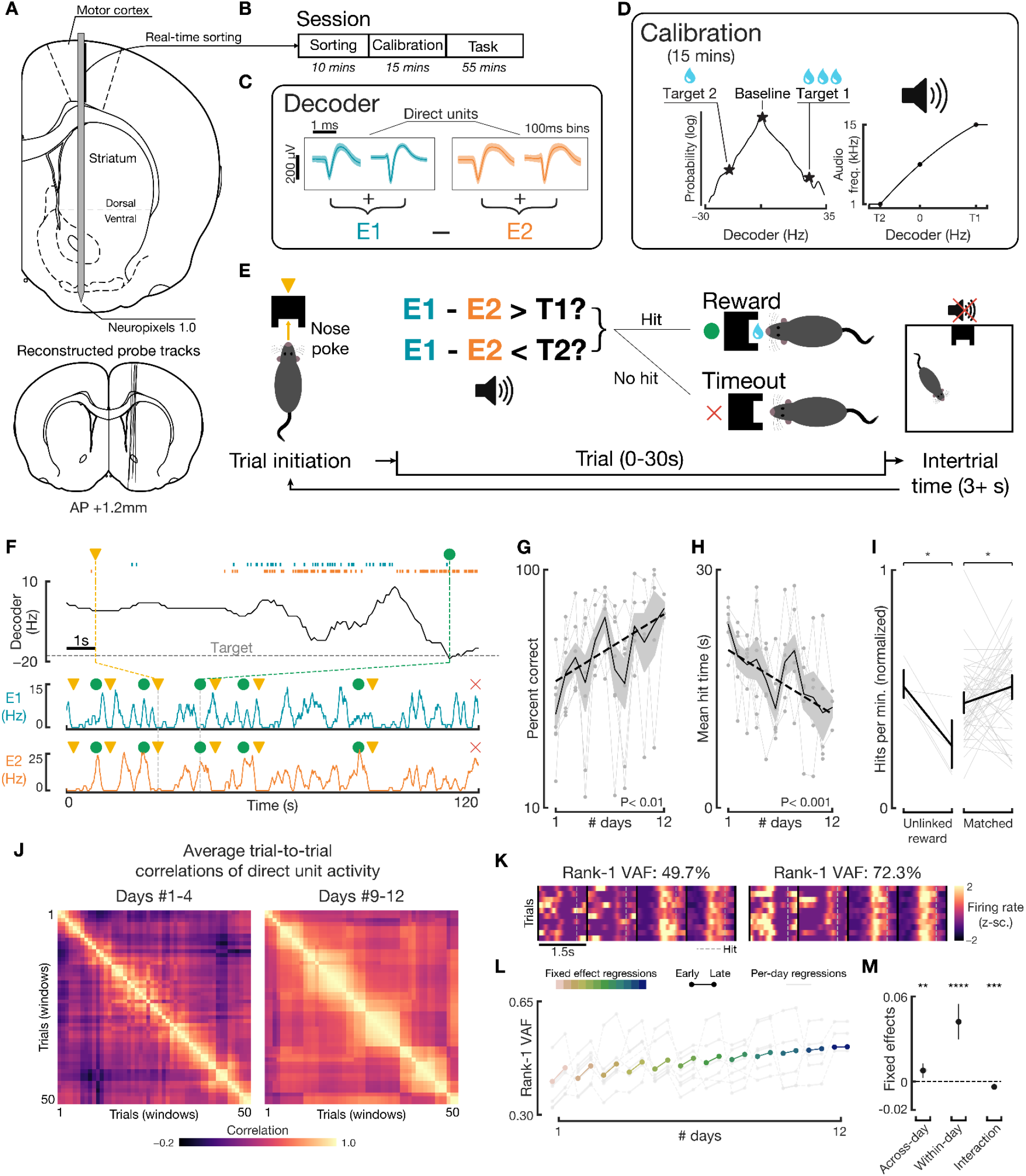
Rats learn a neuroprosthetic task driven by motor cortex units via an increasingly consistent neural strategy. **(A)** Targeted chronic implant trajectory, Neuropixels 1.0 probes in rats (n=7). Bottom: probe tracks via histological imaging confirming desired positioning. **(B)** Phases of daily training. **(C)** Example mean waveforms (shading ± 1 SD) of direct units and decoder calculation. **(D)** Decoder targets and its mapping to auditory tone according to calibration period. **(E)** Task structure. **(F)** Example task data, displaying E1 (middle, blue) and E2 firing rates (bottom, orange) and trial initiation (yellow triangle), hit (green circle), and failure (red X). Top: each dot represents a spike. **(G)** Percentage of correct trials increased (p=0.003). Black line & shading is mean ± 1 SEM; dotted line regression line. **(H)** Mean successful trial length decreased (p=0.0003). **(I)** Hits per minute (max-normalized per-session) decreased after unlinking of reward from the task (left; p=0.02, n=5 sessions), whereas performance-matched sessions increased (right; p=0.04, n=48 sessions). **(J)** Pairwise correlations of trial direct unit activity, averaged across all sessions in the days #1-4 (n=7 rats) and days #9-12 (n=6 rats). **(K)** Rank-1 variance-accounted-for (VAF) computation. **(L)** Regressions from linear mixed model (fixed effects: colored lines; per-day regressions incorporating random effects: gray lines) displaying changes in Rank-1 VAF both within-day (early vs late trial windows, indicated by endpoints of each line) and across-day (x-axis). **(M)** Fixed effects (slopes) from linear mixed model. Across-day slope (Rank-1 VAF per day) 0.008, p=0.006; within-day slope 0.042, p<0.0001; interaction term −0.0038, p=0.0004. For all, bars indicate 95% confidence interval.

Over learning, rats improved in both the proportion of successful trials (p=0.003; Figure 1G) and the average length for successful trials (p=0.0003; Figure 1H). Unlinking reward from the task in a subset of high-performing, late-learning sessions significantly decreased performance, whereas normal sessions with matched initial performance increased in a similar time frame, confirming goal-directed behavior. (Figure 1I). Additionally, while differing reward amounts were provided at the two decoder targets, rats did not prefer targets based on reward amount, direction, or calibration bias, but instead preferred whichever target they initially achieved, a strategy correlated with higher performance and with learning (Supplementary Figure 2). To this end, further analysis in this study considers only the successful trials for the rat’s preferred target on that day (average of 86.8% successful trials per session achieved the same target), unless otherwise noted.

We then investigated rats’ neural strategies by analyzing activity of the direct units surrounding target achievement. Strong trial-to-trial correlations in direct unit activity emerged by late training (Figure 1J), suggesting rats converged to a single consistent strategy each day. Quantifying consistency via principal component analysis (Figure 1K), direct units’ activity increased in consistency both across- and within-day, plateauing in late training (Figure 1K-M). Thus, rats progressively refined rewarded cortical activity patterns, likening such cortical activity to that of skilled movements (*3, 18*). These findings also complement those of previous neuroprosthetic studies (*2, 10, 19*), as we selected direct units along a columnar, rather than the typical horizontal, axis, within which an existing, underlying network structure may have facilitated neuroprosthetic learning (*20, 21*). Hence, this neuroprosthetic task established a suitable framework to study credit assignment to cortical units.

We then asked how individual units in striatum responded throughout this task. Striatal units (n=4633) exhibited widely varied, yet stereotyped, response profiles according to task events. Many striatal units were “reward-responsive” (n=1193/3943 in dorsal striatum, n=361/690 in ventral striatum), characterized by a significant positive or negative modulation in the unit’s mean firing rates in the seconds before and after hit (Figure 2A-G). These units’ firing rates modulated across a wide temporal range. Such modulation was not visible in the same units during simulated hits within the preceding calibration period (Supplementary Figure 4). DS reward-responsive units, on average, modulated both earlier and quicker than those in VS (Figure 2H). Additionally, the relative proportion of reward-responsive units increased over days, with roughly twice the proportion in ventral versus dorsal striatum (Figure 2I). These results indicate a broad heterogeneity among striatal units suggesting different, emergent encodings during the task.

**Figure 2.**
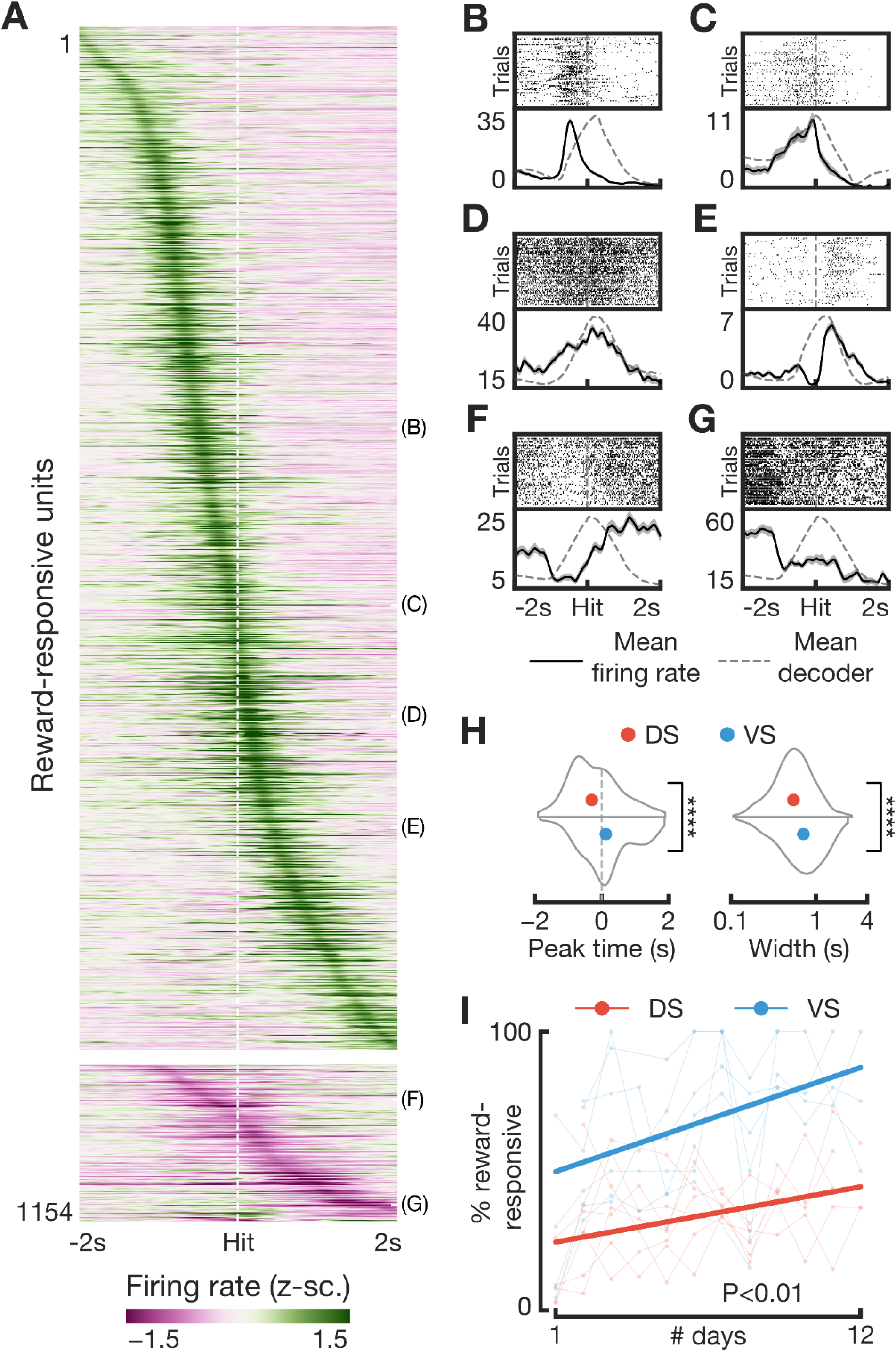
Striatal units develop varied and stereotyped behavior surrounding task hit. **(A)** Mean firing rates (z-scored) of reward-responsive striatal units during the 2 seconds preceding and following a task hit (white dotted line). Top block: up-modulating units; bottom block: down-modulating units. **(B)-(G)** Example reward-responsive striatal units with narrow (B-C-E) or broad (D-F-G) modulations. Top: raster spike plot, time-locked to hit (gray dotted line in center). Bottom: mean firing rates of the same unit (blue; shading 1 SEM) overlaid on mean decoder value (gray dotted line). **(H)** Distribution of peaks (left) and widths (right) of the mean firing rates for all significantly modulating dorsal (red) and ventral (blue) striatal units. Peaks (p<0.0001) and widths (p<0.0001) differ significantly. **(I)** Change in relative proportion of significantly modulating units across days (increase, p=0.004).

In order to ascertain what this stereotyped activity within striatum might encode, we probed the relationship of these reward-responsive striatal units to the task’s two primary means of external feedback: the auditory tone mapped to the decoder output and reward. To quantify changes in striatal representation during particular scenarios deviating from normal task structure (Figure 3A), we compared each unit’s firing rates to the scenario’s preceding trials’ mean firing rates via correlation and subsequent normalization (“Δr”, Figure 3B; Methods).

**Figure 3.**
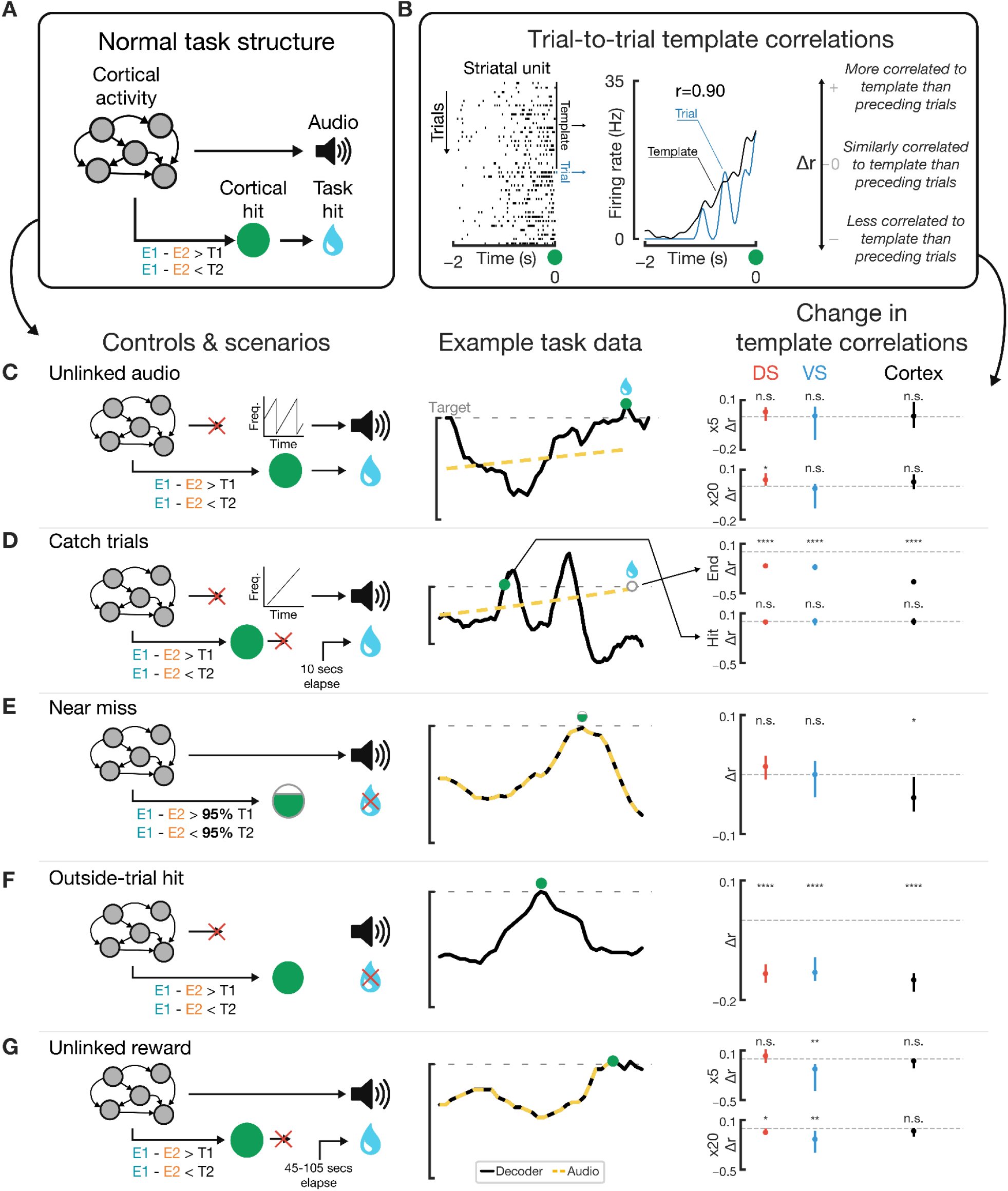
Striatal task-locked activity is independent of external feedback and sensitive to reward availability. **(A)** Normal task structure illustration. **(B)** Trial-wise template correlation (“Δr”) illustration. **(C)** In particular sessions, audio was unlinked from cortical activity. Right: Δr in the 5 (top) and 20 (bottom) trials after unlinking audio. **(D)** Catch trials unlinked both audio and task output. Right, top: Δr for ends of catch trials. Right, bottom: Δr for cortical hits performed during catch trials, despite no reward given. **(E)** Δr for instances at >95% target but no hit. **(F)** Δr for hits outside of a trial, despite no reward. **(G)** In particular sessions, reward was unlinked. Right: Δr in the 5 (top) and 20 (bottom) trials after unlinking reward. For all, n.s. p > 0.05; * p < 0.05, ** p < 0.01, *** p < 0.001, **** p < 0.0001.

In a subset of late, high-performing days (n=7 days), we unlinked the auditory tone from the decoder output in all remaining trials; instead, auditory tone frequencies repeatedly swept a frequency range (Figure 3C). However, reward remained dependent on the decoder output reaching either target. Strikingly, after unlinking the audio, we observed no significant changes in performance (Supplementary Figure 5). Concomitantly, we observed no changes in striatal activity nor in direct units’ activity, with representation even growing more consistent in dorsal striatum after 20 trials (Figure 3C). In addition, we analyzed striatal activity during catch trials, in which the rat’s behavior was entirely decoupled from the task: the auditory tone linearly ramped across the full frequency spectrum over 10 seconds, always ending in a reward regardless of the rat’s neural activity (Figure 3D). At the end of these catch trials when reward was given, striatal activity significantly differed relative to the prior successful trials (subplot “end” in Figure 3D), even for catch trials occurring during earlier or more poorly-performing periods (Supplementary Figure 6). Additionally, despite not resulting in a reward, rats still occasionally performed a cortical hit (i.e., when the decoder output reached a target) during catch trials; striatal activity during these catch-trial hits did not significantly differ (Figure 3D). Taken together, these results demonstrate that striatal activity is not dependent on any external measure of task progress within a trial and does not encode reward anticipation or reward retrieval.

To assess striatal activity’s relationship to reward, we then probed two additional scenarios in which cortical pattern execution did not lead to reward achievement. “Near-miss” cortical activity, i.e. activity that led to decoder output close to, but not sufficient for, achieving a reward, did not lead to significant changes in striatal representation (Figure 3E). Conversely, when a cortical hit that would typically grant reward was executed outside of a trial, we observed significant differences in striatal representation relative to preceding successful trials (Figure 3F), even for those hits where the activity of direct units was more similar to preceding trials (Supplementary Figure 7). Finally, to probe the sensitivity of this striatal representation to devaluation, in a subset of late, high-performing days (n=5 days), we stopped delivering the water reward as a result of cortical hits (Figure 3G). As mentioned above, task performance significantly decreased in the minutes following this unlinking (Figure 1I). Within twenty trials following the unlinking of reward, both dorsal and ventral striatal representations significantly changed. These results show that this striatal representation is dependent on the link between cortical activity and reward.

After interpreting these findings concerning reward and independence from external feedback in the context of a cortically-driven task, we then focused on the reward-related relationship between cortical activity and the striatum. Suggesting a specific focus on the cortical activity driving behavior, we found statistically significant correlations on the shared latent variable between motor cortex and striatum, with direct units’ activity correlated more strongly to striatum than indirect units’ activity at both single-unit (Supplementary Figure 8) and population (Figure 4A) levels.

**Figure 4.**
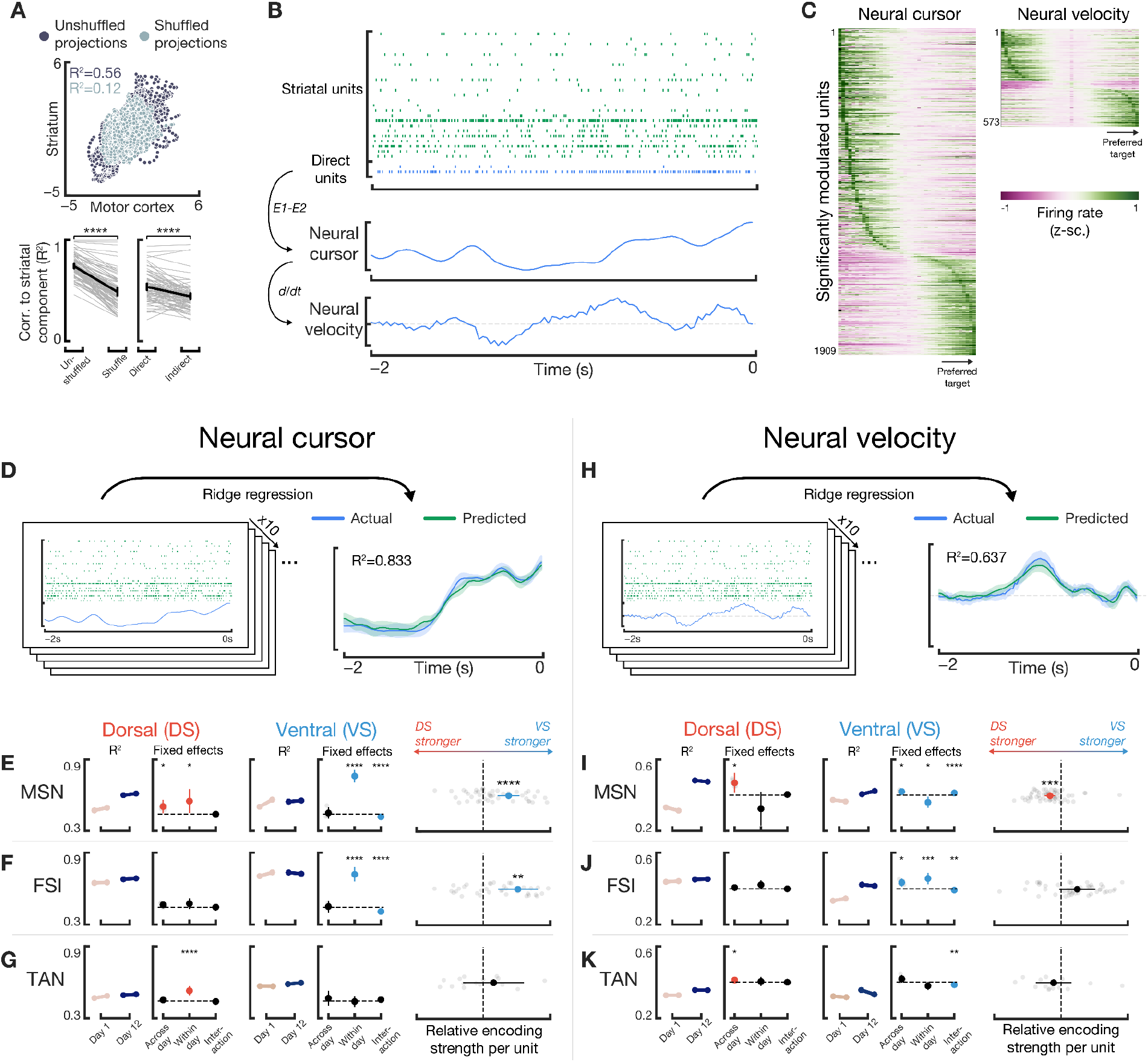
Striatum heterogeneously models both the proximity of cortical activity to reward and the changes of that proximity. **(A)** Top: Cross-area activity between striatum and motor cortex for unshuffled data (light gray) or shuffled data (dark gray). Bottom: coefficient of determination for cross-area activity for each session (light thin lines) or in average (black) when cross-decomposing unshuffled and shuffled motor cortical data (left, p=1e-32 different) with striatal data and when cross-decomposing striatal data with direct units’ data vs indirect units’ data (right; p=2e-11 different). **(B)** Example of the “neural cursor” and its time derivative, “neural velocity” within the 2 seconds preceding cortical hit. **(C)** Tuning to neural cursor (left) or neural velocity (right). Only units deemed significantly modulating are shown. Units are ordered by max bin. **(D)** Population model utilized to analyze encoding strength of the neural cursor. Actual (blue) and predicted (green) cursor values (shading SEM). **(E-G)** For each section: left: regressions from linear mixed model of cross-validated, unit-normalized R^2^ for various subgroups of units on day #1 and day #12. Right: fixed effect slopes (in log-transformed R^2^, see Methods) for neural cursor encoding; positive slopes indicate increases in encoding strength. Faded gray dots indicate random effects, when applicable. Third section: difference between DS vs VS encoding strength per unit recorded. Values farther right indicate a stronger relative encoding for VS than DS; dotted line indicates no difference. Putative cell types for each row are: medium spiny neurons (MSN), fast-spiking interneuron (FSI), and tonically active neurons (TAN). **(H-K)** Same as (D-G) except with neural velocity, instead of neural cursor. For all, * p < 0.05, ** p < 0.01, *** p < 0.001, **** p < 0.0001, otherwise not indicated p > 0.05; bars represent 95% confidence intervals.

We then considered two particular reward-related quantities derived from direct units’ activity as candidates for striatal encoding. First, by utilizing a neuroprosthetic task, we *a priori* define the true relationship between cortical activity and reward via the decoder. Thus, the projection of firing rates of direct units onto this decoder subspace, termed the “neural cursor” henceforth, directly represents the proximity of cortical activity to reward (Figure 4B). Secondly, seeking a continuous analog of action-related measures from more typical motor tasks (*22–24*), we computed the time derivative of the neural cursor, termed “neural velocity” henceforth, to assess encoding of the change in proximity to reward of task-relevant cortical activity (Figure 4B). Many individual striatal units displayed consistent, significant tuning to the neural cursor and neural velocity spectrum (Figure 4C). Seeking to interrogate population-level effects, we then analyzed encoding of neural cursor and velocity within populations of striatal units, taking the cross-validated coefficient of determination (R^2^) from a ridge regression model as a measure of encoding strength (Figure 4D).

Considering all striatal units, we found a strong encoding of the neural cursor within the striatum across all days of training, with a significant increase in encoding strength within each day of training (p=0.0006; Supplementary Figure 9). To investigate specific effects of striatal regions and putative cell-types (Supplementary Figure 10) on the encoding of the neural cursor we repeated population-decoding analysis across different subsets of striatal units. We observed heterogeneous trends dependent on both cell-type and region (Figure 4E-G). In particular, both putative medium spiny neurons (MSNs, Figure 4E) and putative fast-spiking interneurons (FSIs, Figure 4F) in the VS exhibited significant within-day increases in encoding strength (p<1e-10) and stronger encoding than their DS counterparts (p<0.002). Regarding neural velocity, we observed moderate encoding strengths (Figure 4H; Supplementary Figure 9). However, in contrast, we observed strong across-day increases in encoding strength (p<1e-4), with no significant changes within-day (p=0.38; Supplementary Figure 9). Repeating analysis across cell-types and regions (Figure 4I-K), DS MSNs exhibited across-day increases in neural velocity encoding (p=0.025) and significantly stronger encoding than that of their VS counterparts (p=0.0002; Figure 4I). Finally, neural cursor and velocity encodings were independent of auditory feedback but dependent on reward (Supplementary Figure 11) and were stronger than time-related alternative hypotheses (Supplementary Figure 12). Combined together, these data provide evidence that the striatum heterogeneously and simultaneously encodes both the proximity of task-relevant cortical activity to reward and its change over time.

Ventral striatum plays an essential role in enabling stimuli to tune associations between behavior and reward, as VS lesions impair particular forms of learning in instrumental conditioning tasks (*11, 25–27*). However, while prior studies (*16, 28, 29*) congruently show VS firing rates can modulate as animals approach reward, they do not differentiate whether such VS encoding represents stimuli or the rewarded behavior that stimuli might predict. By conditioning a neural pattern, rather than behavior, and subsequently unlinking external stimuli, we provide strong evidence that VS forms an internal representation of the proximity of task-relevant cortical activity to reward, not of stimuli. These results thus shed light on the precise role of VS during learning. Such an internal model may arise from integration across VS’s highly-convergent inputs (*30*) from hippocampus (*31*), amygdala (*27*), and prefrontal cortex (*32*), each likely containing distinct, and perhaps even competing, information (*33*).

Accurate internal encoding of proximity to reward engenders accurate reward-prediction errors critical to learning (*34*). We observe remarkably strong neural cursor encoding in VS, a rapid change in VS representation immediately following task devaluation, and significant within-day increases in neural cursor encoding strength. An accurate, updateable encoding of reward proximity underlies temporal credit assignment in computational models of reinforcement learning (*6*), and thus VS may manage temporally delayed reinforcement in the brain. In particular, via strong bidirectional connections with the ventral tegmental area (VTA) (*10*), this encoding may facilitate appropriate dopamine release at the time of reward across both striatum and cortex, known to be critical for skill learning (*5, 10*).

Shifting focus to dorsal striatum, our stereotyped patterns of dorsomedial striatal units in response to target achievement echo task-locked firing rate modulations observed in prior neuroprosthetic (*9*), navigation (*35*), and motor (*36*) tasks. In conjunction with dorsolateral striatum and dopamine, task-locked dorsomedial activity is hypothesized to guide selection and shaping of actions by coincident activation with cortical activity that leads to the desired action (*36, 37*). The assembly of shorter, behavioral “syllables” into coarser behavioral actions is mediated by the striatum (*38*), implying this striatal influence in goal-directed action selection may be continuous as an action unfolds (*24, 39*).

Our results show DS forms an internal model of the cortical activity responsible for changing the animal’s proximity to reward. In congruence with this theorized role in action selection and shaping, DS may leverage such a model to continuously provide immediate feedback to the motor cortex via downstream thalamic circuits (*32*) to quickly positively or negatively bias unfolding cortical activity. Critically, a mapping of the neural velocity would provide more immediate feedback relative to that of the neural cursor, reminiscent of the responsiveness benefits of derivative-based controllers widely utilized in control theory (*40*). Such a mechanism can then underlie structural credit assignment: by dynamically filtering cortical activity into desired and undesired activity – i.e. activity bringing the animal closer and farther from reward, respectively – dorsomedial striatum can participate in a positive-feedback recurrent loop biasing those cortical neurons leading towards reward (*41*). DS’s involvement in structural credit assignment may primarily manifest during goal-directed learning, before DS becomes less engaged (*12, 35*) and less sensitive to task context (*42*) as behavior becomes more automated.

Our results demonstrate an internal, continuous model of the proximity of cortical activity to reward within the striatum, with differential representation across dorsal/ventral region and cell types. Such a model connects cortical performance to the neural circuits underlying goal-directed learning, an instrumental pathway to solve the credit assignment problem in the brain.

## Acknowledgements

We thank V. Athalye and I. Rodriguez-Vaz for thoughtful discussion and G. Shvartsman for assistance with histological procedures.

## Authors’ contributions

PB, NVL, RMC and JMC contributed to the conceptualization of the manuscript. PB performed the experiments and curated the data. PB and NVL developed the software and performed the analysis. RMC and JMC supervised the investigation. PB and NVL wrote the manuscript and created the visualizations with considerable contributions from RMC and JMC.

## Materials and Methods

### Animals

All rat experiments were performed in compliance with the regulations of the Animal Care and Use Committee at the University of California, Berkeley. Singly housed, male Long-Evans rats on a 12h light/dark cycle weighing 200-300 g were used for the experiments. All rats that had at least 4 well-isolated units in the motor cortex performed the neuroprosthetic task for as many days as possible, until no more motor cortex units remained or until sufficient days of training elapsed. Only rats (n=7) that performed at least 8 days of the task were included in analysis. In 1 rat, 2 days in late training (days #14 and #17) were omitted from analysis due to a different neuroprosthetic task structure attempted. Otherwise, no rats, trials, or sessions were excluded from analysis.

### Implant Construction

All rats were chronically implanted with Neuropixels 1.0 probes, mounted within a custom-designed, lightweight 3D-printed enclosure containing both the probe and the Neuropixels headstage PCB. This enclosure added onto a prior study’s design (*43*) and enabled quick and stress-free probe connections by chronically mounting the headstage and utilizing snap-fit joints to connect the headstage PCB. A transport-friendly container utilized for safekeeping and for sterilization was designed to secure the probe mounted within its enclosure. Explantation of probes was possible with this implant. Construction was largely similar to that of Luo et. al (*43*).

### Rat Surgery

Targeted stereotactic coordinates were 1.4mm anterior to bregma and 2mm lateral from midline, though exact implant coordinates varied based on cortical vasculature and removal of dura mater. Probes were lowered as deep as possible, up until approximately 8.5mm deep from the cortical surface.

Fully assembled probes within their enclosures were sterilized with ethylene oxide gas sterilization. Immediately before surgery, probes were maximally lowered into an excess quantity of lipophilic dye for histological probe tracking (Vybrant DiI Cell-Labeling Solution, ThermoFisher Scientific, Waltham, MA) repeatedly for several seconds.

After approximately 1 week of manual handling, rats of age 8-9 weeks old, 200-300g were anesthetized with isoflurane (0.5-3%), set in a stereotactic frame (Kopf, Tujunga, CA, USA). Artificial tears were applied, and a closed-loop heating pad utilized to maintain 35°C (RightTemp, Kent Scientific Corporation, Torrington, CT). Dexamethasone (0.75 mg/kg) and buprenorphine (0.05 mg/kg) were administered subcutaneously for anti-inflammation and analgesic properties. Saline (10 ml/kg) was administered subcutaneously for hydration. Bupivacaine (1.5 mg/kg, diluted further by 50% with saline) was administered locally to the surgical site to provide secondary local anesthesia. Hair was cleared with an electric shaver and fully removed with brief application of hair-removal product. After verifying surgical levels of anesthesia, rats were secured in blunted ear bars, and the surgical site was sterilized with isopropyl alcohol and chlorhexidine. A midline incision was then made with a scalpel, and skin retracted with Alm retractors (Fine Scientific Tools, Foster City, CA). A spatula and forceps were used to clean the skull of all overlying tissue while keeping the skull moist with sterile saline kept on ice. Additionally, hydrogen peroxide was used to aid removal of remaining tissue and to aid in drying the skull. A #15 blade was used to gently yet firmly scrape the surface of the skull in a cross-hatch pattern to increase roughness of the skull for improved eventual adhesion of dental cement. The skull was fully dried and all bleeding contained.

6-8 1mm diameter holes were drilled around the perimeter of the skull utilizing a custom-made impedance-sensing automated drill system which stopped precisely at the detection of dura mater and/or CSF, based loosely on the system of another study (*44*). M1×2mm screws (McMaster Carr, Robbinsville, NJ) were then firmly screwed into each hole until dura was just touched. A rectangular craniotomy of 2mm x 2.5mm centered around the target implant site was then drilled using the same automated drill system, which enabled manual specifications of craniotomy depths and shapes. The perimeter of the craniotomy was repeatedly drilled until the piece of skull could be excised gently with minimal pressure. During excision of skull, constant irrigation with cold saline minimized sticking of dura to excised skull pieces. Once the dura mater was exposed, a 30G needle with its tip bent at 45 degrees was utilized in conjunction with angled forceps (Dumont #5/45 forceps, Fine Scientific Tools) to very gently cut and remove patches of dura mater in areas clear of large vasculature. Bleeding frequently occurred despite utmost caution; in these cases, gelfoam soaked with cold saline was gently moved over bleeding sites until bleeding stopped. Only the minimal dura was removed to enable implantation of the probe clear of surface vasculature. A piece of saline-soaked gelfoam was then placed in the craniotomy space, and saline periodically applied as necessary to ensure the exposed cortex was hydrated.

The skull was then redried completely. The probe enclosure was then mounted stereotactically (Model 1766-AP Cannula Holder, Kopf) and straightened appropriately. The enclosure was moved near to its desired implant site, and the ground wire was connected by repeated wrapping around 2-3 skull screws. Ground wire/screw connectivity was confirmed with impedance tests between the screw and the rat’s paw. After re-ensuring the skull was completely dry, C&B Metabond (Parkell, Edgewood, NY) was used to completely cover the exposed skull, including all skull screws and the ground wire. After the cement dried, gelfoam within the craniotomy was removed, and the probe tip was lowered to the cortical surface using a 25x zoom surgical microscope. Utilizing custom-built software controlling a vertically-mounted linear translation stage (ThorLabs NRT100/M, Newton, NJ) mounted on top of passive vibration isolation legs to minimize vibration (ThorLabs PWA075), the probe was then carefully and slowly lowered to minimize damage to brain tissue and minimize eventual glial encapsulation of the probe. The first 1mm of insertion was lowered at 1-2 microns per second, the next 5mm of insertion was lowered at 4 microns per second, and the remaining insertion was performed at 1-2 microns per second; saline was periodically applied to keep cortex hydrated, and the surgical table was minimally touched to reduce induced vibrations. When the probe was within 1mm of its final depth, a small quantity (~1-2 microliters) of soft silicone elastomer (DOWSIL 3-4680, Dow) was carefully injected with a micropipette to seal the craniotomy while minimizing vibrations to the surgical table; too much elastomer could lead to overflow onto the skull and prevent proper sealing with cement in subsequent steps. The probe was then lowered to its final, maximal depth (<= 8.5mm or just before the implant touched the skull). We then applied a viscous dental composite (Absolute Dentin, Parkell) to seal the craniotomy and fix the implant in place to the skull. Once the viscous cement completely dried (~4 minutes), less viscous dental cement (Ortho-Jet, Lang Dental, Wheeling, IL) was applied liberally around the implant and skull; the loose skin surrounding the implant was used to gently mold dental cement in order to ensure smoothness and improve surgical time. Rats were given meloxicam (2 mg/kg), another supplement of dexamethasone (0.2 mg/kg), and saline (10 mL/kg) subcutaneously, and antibiotics were liberally applied. Rats received a taper schedule of dexamethasone (1.0 mg/kg 2 days, 0.5 mg/kg 1 day post-surgery), and were allowed five days post-surgery to recover before experiments began. Antibiotics were applied daily; rats that did not recover weight by five days post-surgery were allowed extra time to recover as needed.

### Electrophysiology

Unit activity from Neuropixels 1.0 probes (*17*) was recorded and visualized via OpenEphys GUI (*45*) (Open Ephys), and all neural data was streamed to disk. Custom software written in Python and C++, spanning both standalone services and OpenEphys plugins, enabled real-time readout of spikes of selected units. Services were networked with a high-throughput, low-latency streaming framework with C++ and Python bindings, River (*46*).

Each day, 2 2-minute baseline sessions were first recorded in order to determine the enabled channels for the Neuropixels 1.0 probe; one session captured the deepest 384 channels, and another session captured the next-dorsal block of 384 channels. We then utilized Kilosort2 (www.github.com/MouseLand/Kilosort2) to sort each session individually, from which the 384 channels capturing the most units across the probe length were selected via an optimization algorithm (*47*). A third 2-minute baseline session was then performed with this final channel configuration and was sorted via Kilosort2.

Utilizing a custom-built GUI, 4 well-isolated units within the motor cortex were selected from these Kilosort2 results as “direct units”, with preference given to selecting units with similar waveforms, depths, and/or firing properties as previously-selected direct units for that rat. In cases where similar units were not obviously available, direct units were chosen from similar depths within the motor cortex as preceding days. All rats exhibited a gap in well-isolated units beginning between 2.0mm and 2.5mm from the cortical surface, putatively corresponding to the dorsal edge of white matter tracts of the corpus callosum; direct units were chosen strictly dorsal to the start of this gap. Note that, contrary to previous neuroprosthetic studies where the use of multielectrode arrays meant direct units distributed horizontally while at similar depths, here the use of a Neuropixels depth probe means direct units distributed along a dorsal/ventral axis, within approximately the same anterior-posterior/medio-lateral position. Thus, direct units existed between approximately 1mm and 2mm deep from the cortical surface; the average difference between the two farthest direct units in any given session was 0.5mm, and the average depth of a direct unit was 1.4mm relative to the cortical surface. Post-mortem histology (described below) confirmed the locations of direct units within the motor cortex.

We then desired to adopt Kilosort2-identified spikes for our 4 selected direct units into a methodology commonly used for online spike sorting: classification via ellipsoids within a PCA-determined space of waveforms of threshold crossings. To do this, for each selected direct unit, we identified an appropriate threshold for each relevant channel based on that channel’s RMS noise in order to perform threshold-crossing extraction. From these threshold crossings, we then identified and subsequently projected these voltage waveforms into a 3-dimensional space via PCA. Particular threshold crossings corresponding to Kilosort2-identified spikes then formed the putative “ground truth” for online sorting, and a best-estimate ellipsoid in PC-space was constructed via stochastic gradient descent to maximize accuracy (*48*). The parameters for the PCA projection and subsequent ellipsoid classification were then transferred to OpenEphys for online spike identification. For subsequent sessions that required real-time online sorting, an OpenEphys program was used that processed data online in similar fashion to Kilosort2: common median referencing across all channels, followed by a 150 Hz, 3rd order highpass filter. Online-identified spikes for each of the 4 direct units were then streamed out from OpenEphys in real-time via River. Firing rates were then continuously computed via these streamed spikes in 100ms bins and smoothed using a running average of 10 bins (1 second). Frequencies for the auditory tone were simultaneously computed as well, according to the thresholds set during the calibration period, described below.

In a subset of rats (n=4), improvements were enacted to increase the agreement between online-identified spikes and Kilosort2-identified spikes for the (putatively) same unit within a session, in cases where neural recordings were not fully stationary within a day. Though not common, some direct units tended to appear to slowly drift along the dorsal/ventral axis, i.e. along the length of the probe. To combat this, instead of a single-channel’s 3 dimensions in PC space, each unit was identified using up to a 15-dimension ellipsoid, with 3 PC dimensions originating from each of 5 channels stacked along the dorsal/ventral axis. Then, every 6 minutes with a 3-minute overlapping window, PC scores of all threshold crossings on a unit’s channels were fit with a Gaussian mixture model (sklearn.mixture.GaussianMixture) with a number of components chosen to minimize the Akaike Information Criterion (AIC). The cluster containing the highest proportion of online-identified spikes became the new “ground truth” for that unit’s online sorting, and a new best-estimate ellipsoid was fit to this cluster and formed the new basis for identifying spikes for that given unit online. This updated clustering was only accepted if it had at least 70% agreement with the prior sorting. This methodology thus enabled tracking of slow changes in neural recordings across channels.

### Behavioral Task

Upon recovery from surgery, rats were deprived of water for 24 hours before initiation of experiments. During training, rats only received access to water during the behavioral task, unless supplemental water was needed to maintain 90% body weight. After initial sorting and selection of direct units were performed as described above, rats had a 15-minute calibration period, in which they freely moved around the cage and were passively given water rewards paired with a reward tone every 45-105 seconds (uniformly random). After 15 minutes, the calibration period data was used to determine assignments of the 4 direct units into ensemble #1 (E1) and #2 (E2), the positive (T1) and negative (T2) neuroprosthetic targets, a baseline value, and the mapping of decoder output to the frequency of the auditory tone. The goal of calibration was to find the set of parameters leading to a success rate during calibration of 35-40% of trials.

In order to set these parameters, we simulated the task performance of the rat during this 15-minute calibration period for all 6 combinations of E1 and E2 and across a sweep of thresholds for T1 and T2. In particular, T1 and T2 values were selected based on exhaustive search that minimized the difference between simulated and goal rates of success (35-40%). E1 and E2 assignments were then selected as the assignments that yielded a success rate close to the goal, had subjectively fair thresholds, and were balanced between T1 and T2 rewards. The baseline was set as the mean E1 - E2 value during calibration. The auditory tone’s frequency was set according to a 2nd-order polynomial fit interpolating decoder values of T2, 0, and T1 to frequencies 1, 8, and 15 kHz, respectively. These frequencies fall within the auditory spectrum of the rat. The task structure used during simulations was identical to the true task structure, and was as follows (and as illustrated at a high-level in Figure 1).

First, firing rates were first computed using spike counts in 100ms bins and smoothed with a 1s rolling average filter. Then, for each time bin, decoder output was computed as

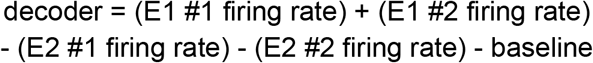

Note that, elsewhere in this study, we might refer to the decoder as *E1 - E2* as shorthand for the above full formula for the decoder. Trials were initiated once the decoder output crossed zero at least once between trials and once at least 1 nose-poke was detected between trials. During a trial, if the decoder output exceeded either T1 or T2 for 1 bin, the trial was declared successful; an extended tone with frequency matching that target’s frequency was played, and reward was delivered simultaneously. Reward was 30% sucrose water. Each rat was randomly assigned a target to be more highly rewarded, where rewards for this target yielded three times as much water as the other target; this assignment was held constant for the duration of experiments for that rat. If 30 seconds elapsed after trial initiation without decoder output exceeding either T1 or T2, the trial was declared failed, and a white noise sound played. After either reward delivery after a success or a white noise tone after a failure, an inter-trial period of at least 3 seconds was enforced, after which the animal was free to initiate another trial. The auditory tone played continuously during a trial and was muted when no trial was ongoing. Every 20 trials (regardless of success or failure in those 20 trials), catch trials were performed, in which the auditory tone’s frequency either ramped from 1 to 15 kHz or 15 to 1 kHz over 10 seconds regardless of the rat’s direct unit activity, after which the animal would hear a reward tone and receive a reward (Figure 3D). Each day’s session would end after approximately 55 minutes of task time or if the animal reached satiety and stopped initiating trials.

Additionally, in a subset of high-performing (>= 75% trial success rate in a trailing 10 minute window and <= 6 mL water consumed), late-learning (day >= 8) sessions, we unlinked either the auditory tone (Figure 3C) or the reward (Figure 3G) from cortical activity for the remainder of that day’s trials. In the case of the former, linear sweeps of frequencies from the lowest (1 kHz) to the highest (15 kHz) ends of the spectrum were performed over a period of 10 seconds and repeated; otherwise, the task structure remained identical. In the case of unlinking reward, we instead delivered a reward every 45-105 seconds, approximately matching the rat’s calibration success rate (~35-40%); otherwise, the task structure remained identical.

### Explantation & Histology

After the completion of experiments, rats were first anesthetized with isoflurane in preparation for explantation. After surgical anesthetic levels were confirmed, rats were fixed in a stereotactic frame. Screws securing pieces of the 3D-printed implant were loosened, and the headstage PCB was removed and disconnected. After ensuring vertical alignment to minimize damage to the probe, the same motorized system used to implant the probe was used to gently remove the probe from the implant and brain. Upon successful explantation, the probe was immediately soaked in 10% Tergazyme solution for 24 hours. After this, the probe was soaked in deionized water for 5 minutes, and then soaked in isopropyl alcohol for 1 minute. In cases where residual tissue or elastomer remained on the probe, the probe was cleaned via an ultrasonic cleaner (CREWORKS) filled with isopropyl alcohol, as residual matter would interfere with subsequent recordings. For stuck-on elastomer, the probes were soaked in elastomer solvent (DS-2025, Dow) for 18-24 hours, followed by a 10-minute soak in DI water and then another minute in an ultrasonic cleaner with isopropyl alcohol.

Immediately after explantation of the probe, rats were injected with sodium pentobarbital and transcardially perfused with PBS followed by 4% paraformaldehyde. Brains were removed and post-fixed in 4% paraformaldehyde overnight at 4°C, after which they were stored in PBS at 4°C. When ready, brains were then transferred to 15% sucrose/PBS solution until they sunk, and then a 30% sucrose/PBS solution until they sunk. Brains were then mounted and sliced via a cryostat into coronal slices. Brain slices were then serially imaged with DAPI and DsRed. Images were preprocessed to adjust for orientation and contrast via manual scripts. Images were then aligned in three dimensions to the Waxholm Space Rat Atlas (*49*) via the QuickNII tool (*50*). Probe tracks within each slice were programmatically segmented via ILastik (*51*), 3D coordinates of probe tracks spanning each brain’s slices were extracted via Nutil (*52*), and linear probe tracks were finally reconstructed from these 3D point clouds via custom software. 1 rat’s brain was unable to be imaged successfully due to technical issues during the slicing process.

### Analysis & Statistics

Analyses were performed in Python 3.8 (https://www.python.org/) with custom-written scripts utilizing publicly available software packages, including numpy (*53*), scipy (*54*), pandas (*55*), and scikit-learn (*56*). Data pipelines were constructed utilizing Apache Airflow (https://airflow.apache.org/), and the majority of data files were stored in the Parquet file format (https://parquet.apache.org/), a cross-platform, high-performance columnar data format.

Unless explicitly noted otherwise, in all figures, n.s. p > 0.05; * p < 0.05, ** p < 0.01, *** p < 0.001, **** p < 0.0001.

Additionally, where indicated for use below, linear mixed models (LMMs) were implemented via the pymer4 package (*57*). Each model was inspected for homoscedasticity in residuals via the Het-White test or visually. Normality of residuals was not strictly enforced, as LMMs are relatively robust to non-normality in residuals (*58*); instead, kurtosis and skew of residuals of all models were both confined to be less than 3.0 and are stated as needed per model below. Confidence intervals were computed via Wald estimates.

### Behavioral Analysis

Behavioral analysis across sessions focused on percent of successfully completed trials (“% correct”, Figure 1) and the average length of successful trials (“Mean hit time”, Figure 1), considering trials of all targets. Analysis was done with linear regression (Figure 1). To study within-session behavioral changes after the unlinking of reward (Figure 1I) and unlinking of audio (Supplementary Figure 7), we utilized the number of hits per minute, a measure that incorporated both the length of trials and the time the rat took to initiate trials. We took the mean hits/min in the 10 minutes preceding and the 10 minutes following the start of the first unlinked trial; statistical significance was tested via a paired t-test.

Within each day, the rat preferred one target more than the other (Supplementary Figure 2). The preferred target for that day was considered to be the target that had more successful trials for that day. Unless explicitly noted otherwise, all analysis described below utilizes successful trials to the rat’s preferred target for that day.

### Data Analysis: Preprocessing into Firing Rates

Each day’s session was first sorted via Kilosort2. All unit waveforms were manually curated via Phy; units subjectively determined to be noise were excluded. Of these non-noise units, only high-quality units were then utilized for the remainder of this study’s analysis, as defined by: having a mean firing rate of greater than 0.2 Hz both globally and within the 4 seconds centered on the end of each successful trial, having less than 50% ISI violations, less than 50% missing spikes due to low SNR, and a trough in the unit’s mean waveform occurring temporally before its peak to exclude axonal spikes (*59*). Additionally, in 1 rat, during surgery it was noted the probe was angled relative to the brain during insertion, and histological analysis confirmed the ventral portion of the probe did not intersect with the ventral striatum; all units in this ventral portion of this particular probe were excluded from analysis. Finally, the units and spikes identified online as direct unit activity were merged with these Kilosort2-determined units and spikes. For the remainder of the analysis, all “direct unit activity” corresponds to the units/spikes determined online.

Spike counts for each unit were binned at 20ms resolution and divided by bin size to yield (unsmoothed) firing rates. These firing rates were then extracted time-locked to the end of each trial. Owing to the broader binning/smoothing used online (100ms bins with 1s rolling average) compared to that of the offline analysis done here – as well as to non-zero jitter in the experimental setup itself – neural data varied in its degree of alignment with the online-determined “trial end” across trials. In order to correct for this, time shifting on each trial was performed to optimize the trial-to-trial correlations of direct unit activity with the successful trials of each target (*60*) (Supplementary Figure 3). The computed optimal shift for the direct units was then applied to the firing rates of all units to globally align all units to the time of cortical hit. Notably, we only shifted units in time but did not warp time, and the same shift was applied to all units for each trial. Additionally, the shifts per session were adjusted such that the correlations of the unaligned and aligned decoder outputs were maximal, thereby minimizing deviations between the two while increasing overall trial-to-trial alignment. Finally, these aligned firing rates were smoothed with a centered Gaussian kernel with standard deviation 60ms to yield firing rates for each unit. For all subsequent neural data analysis, unless otherwise noted, we utilized these aligned, Gaussian-smoothed firing rates.

Units were then segmented into approximate brain regions according to probe insertion depth. As stated above, all rats exhibited a gap in units beginning between 2.0mm and 2.5mm relative to the cortical surface. Thus, a given unit was considered to be in the dorsal striatum if the channel containing the largest amplitude in the unit’s mean waveform was between 2.5mm and 5.5mm below the approximated cortical surface; all units more ventral than 5.5mm were considered ventral striatal units, and those dorsal to the gap were considered motor cortex units.

Furthermore, the direct units and spikes identified online were duplicated in the motor cortex units and spikes identified by offline Kilosort2 sorting. To identify these offline-sorted units corresponding to the online-sorted direct units, we computed the fraction of spikes between each offline and each online-sorted direct unit that occurred within 1ms of one another. Those offline units that had at least 50% agreement with a direct unit and were less than 120μm away from that direct unit were identified as a duplicated direct unit; these relatively relaxed criteria aimed to establish an upper bound on duplicated direct units. These offline-identified, duplicated direct units were then excluded from Granger causality and cross-correlation analysis of “indirect” motor cortical units in Supplementary Figure 8 and Figure 4, respectively, described below.

### Data Analysis: Direct units

To analyze direct unit activity (Figure 1J-M), firing rates of each direct unit were first z-scored, based on the mean and standard deviations computed from all successful trials towards the preferred target between 1.2 seconds before the hit and 0.3 seconds after (i.e. 75 time bins each trial). Z-scored direct unit activity was then concatenated within-trial across the four units, yielding 300 time bins per trial. Then, we constructed overlapping, sliding windows of 10 trials each, and used 1-component PCA across trials to yield a 300-element projection representing the normalized “average” of the direct unit activity within that window. In Figure 1J, the projections between sliding windows were compared pairwise with Pearson correlation. In Figures 1K-M, we took the variance-accounted-for by this 1-component PCA (Rank-1 VAF) as a measure of consistency of direct unit activity within each window. Note that for this measure and window size of 10 trials, a standard normal Gaussian noise process would expect a Rank-1 VAF of 0.133 (simulations not shown). Sessions with fewer than 10 successful trials were ignored. Additionally, since across-day changes were a focus, only sessions within the first 12 days were considered to ensure at least 3 rats’ sessions were included for each day.

We then utilized a linear mixed model to analyze the changes in Rank-1 VAF over within- and across-day timescales. First, we designated “early trials” as the first 15 windows of 10 trials (i.e. spanning the first 25 trials), and “late trials” as all other windows of trials. Then, we constructed a model with: fixed effects of number of days, indicator of early or late trials (−1 or 1), and an interaction term, a random intercept per session, and a random across-day slope per rat. The model converged and met appropriate statistical assumptions (heteroscedasticity test p=0.28, kurtosis/skew of residuals 0.74/0.40).

### Data Analysis: Reward-responsive modulation

To analyze striatal unit activity, each unit in the striatum was first z-scored, based on the mean and standard deviations of the time periods between 3 and 6 seconds both before the hit and after the hit. Note these windows were intentionally selected outside of the rewarded period to be analyzed in order to capture each unit’s relative baseline. Mean z-scored firing rates were then computed for the period between 2 seconds before and 2 seconds after the hit (i.e. 200 time bins total), and modulation depth for each unit was computed as the difference between maximum and minimum z-scores.

Reward-responsive striatal units (Figure 2) were then determined using a sliding z-score threshold on modulation depth in order to account for varying numbers of successful trials each day. In particular, we computed the z-score threshold corresponding to a 95% confidence that the modulation depth in the unit’s mean firing rates was not generated by a stationary, independent Gaussian noise process with 200 samples per trial. This threshold was the minimum z-score threshold T such that, for a given number of trials N, the following probability was below 5%:

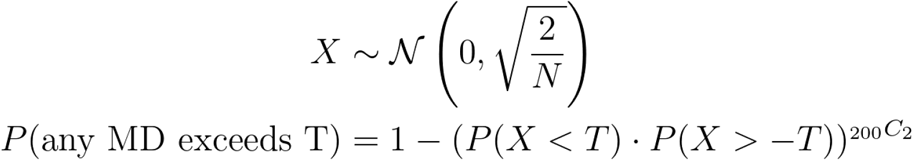

We then ensured all z-score thresholds on modulation depth were non-trivial by setting a minimum threshold of 0.75. This process yielded z-score thresholds such as 1.5 for sessions with 20 trials or 0.75 for sessions with 79 trials or more.

After filtering units according to this z-score threshold, we then identified units as positively modulating or negatively modulating depending on whether the magnitude of the max z-score or min z-score were greater. The time at which this peak occurred was denoted as the peak time (Figure 2H). Then, to further reduce the effects of noise or trial size on determination of reward responsiveness, only those units also exhibiting a mean firing rate at the peak time significantly different than zero were finally deemed reward-responsive (Wilcoxon signed-rank test, p-value < 0.05). The width of the modulation (Figure 2I) was determined according to 50% of the peak’s prominence in the mean firing rate (scipy.signal.find_peaks with rel_height=0.5). Statistical tests on the distributions of peak time and modulation width used medians (testing different than zero) or the nonparametric Mann–Whitney U test (comparing distributions). Changes in relative proportions of DS and VS reward-responsive units were modeled with a linear regression simultaneously including both DS-specific and VS-specific slopes and intercepts.

### Data Analysis: Manipulations

We then sought to analyze changes in activity in reward-responsive striatal units during particular deviations from the normal task structure (Figure 3). In particular, we examined: the ends of each trial within 5 or 20 successful trials after unlinking the auditory tone (Figure 3C) and reward (Figure 3G) from cortical activity; the first cortical hit occurring during each catch trial (if any), as well as the ends of all catch trials (Figure 3D); instances where the local maximum of the decoder output reached 95% of either target T1 or T2 (Figure 3E); and instances where a cortical hit occurred at least 10 seconds outside of any trial (Figure 3F).

For each of these scenarios, we extracted the 2 seconds leading up to a particular scenario (e.g. the 2 seconds preceding a cortical hit). For near misses, outside-trial cortical hits, and catch trial cortical hits, firing rates were first re-aligned according to direct unit activity to include these new instances of cortical hits, in addition to including all successful trials to the preferred target as done initially (Supplementary Figure 3). Then, for each scenario, the mean firing rate for each unit for all preceding successful trials was computed and labeled the “template” for that particular scenario and unit. The firing rate for that scenario was then compared to the template via Pearson correlation. This process was then repeated for the 10 successful trials preceding this scenario, effectively computing a “baseline” correlation of successful trials to their templates relative to this unit and scenario. Δr was then computed for each unit and scenario as the difference between the scenario’s Pearson correlation coefficient and the mean of these 10 “baseline” correlation coefficients. Comparing Δr values across units, instead of raw Pearson correlation coefficients, thus accounted for local differences in stability of the running mean firing rate for each unit. Finally, to analyze these trial-to-trial template correlations, the median Δr value for each unit was taken across scenarios, and then BCa-corrected confidence intervals and p-values were computed using bootstrapping to test if median Δr values significantly differed from zero.

Using this methodology, significant positive values in Δr indicated scenario firing rates closer to the template than preceding trials, i.e. indicating a strengthening representation; Δr values not significantly different from zero indicated a representation not significantly different from preceding trials; Δr values significantly negative indicate scenario firing rates that are farther from the template than preceding trials, indicating a changing representation (Figure 3B).

Trial-to-trial template correlations were computed similarly for cortical activity as for striatal activity, except: the time period extracted was between 1.2 seconds before the scenario and 0.3 seconds after the scenario, firing rates for each direct unit were first z-scored and then concatenated within-trial across the direct units before the computation of correlation coefficients, and statistical tests were computed directly on the Δr values without taking the median across scenarios.

### Data Analysis: CCA

To assess the relationship between cortical and striatal populations, we utilized canonical correlation analysis (CCA, sklearn.cross_decomposition.CCA) to compute the projections that were maximally correlated between the two populations. Similar to principal component analysis (PCA), CCA also aims to reduce the dimensionality of data by finding projections that explain the originating dataset well; however, unlike PCA, CCA finds two components that simultaneously reduce dimensionality across two populations and is thus well-suited to describe cross-area dynamics in neural data (*61*).

To this end, CCA was performed on the firing rates within the trailing 2 seconds of all successful, preferred-target trials, concatenated together. For all below analysis, 10-fold, shuffled cross-validation (sklearn.model_selection.KFold) was used to compute the cross-validated squared correlation coefficient (R^2^) as the mean test R^2^ over the 10 folds.

First, to determine whether there was significant motor cortex / striatum cross-area activity, for each session, we shuffled cortical data in time 100 times, and took the mean cross-validated R^2^ as a baseline. Comparisons were made between this and the unshuffled data via a paired t-test (Figure 4A, left).

Secondly, we sought to compare the specificity of striatal encoding by comparing the population components of striatum and direct units versus the population components of striatum and indirect units (*62, 63*). CCA was performed in two ways: first, cross-decomposing the activity of all striatal units and the 4 direct units for each session, and then cross-decomposing the activity of all striatal units and indirect cortical units. To correct for cortical neuron sample size in the latter, 4 random indirect units were chosen for analysis and the cross-decomposition performed, and this process was repeated 100 times and results averaged. Again, comparisons between striatum/direct and striatum/indirect cross-area correlations were performed via a paired t-test (Figure 4A, right).

### Data Analysis: Granger Causality

Similar to the CCA analysis, we desired to investigate the level of functional connectivity between motor cortex and striatum, but now at the level of individual units.To this end, we computed the directed functional connectivity between all pairs of motor cortex and striatal units via Granger causality analysis (up to 100ms lags allowed, i.e. 5 bins). To establish whether a given value was significant, each unit’s data was shuffled in time 100 times, and a given unit-pair was considered connected if its (unshuffled) F-test value exceeded the 95th percentile of the shuffled F-test values. Un-smoothed neuronal activity was used for this analysis. The same window of firing rate data was used as in the above CCA analysis (i.e. trailing 2 seconds of all successful, preferred-target trials).

### Data Analysis: Neural cursor and velocity computation

Seeking concrete, task-relevant, cortically-derived quantities that could be encoded in the striatal population, we next defined the neural cursor and neural velocity. The neural cursor was defined in the same manner as the decoder utilized for the task: the sum of the firing rates of E1 minus the sum of the firing rates of E2 minus the baseline value computed during calibration. Notably, this differed from the decoder output only in the smoothing parameters: while the decoder was computed with 100ms bins and smoothed causally over 1 second, the neural cursor was computed using the binning (20ms bins) and smoothing (centered Gaussian filter with 60ms standard deviation) used for all analysis. Additionally, since it was computed offline, the neural cursor exactly aligned with binning utilized for all striatal activity. This neural cursor value was finally normalized to values between −1 and 1, where 1 represented the neural cursor value sufficient to hit the preferred target and −1 the other target. The neural velocity was then computed as the time derivative of the (unsmoothed) neural cursor followed by a more broad smoothing (centered Gaussian with 120ms standard deviation) than the cursor, since time differentiation significantly amplifies noise.

### Data Analysis: Single-unit tuning curves

To construct mean firing rates of individual units at particular points in the neural cursor and velocity spectrum (Figure 4C), we considered all task trial data, across the full length of all trials (though excluding any catch trials and those where reward or audio was unlinked). We first divided the neural cursor spectrum into 40 bins between −1 and 1, and the neural velocity spectrum into 40 bins between −3 and 3. Then, each unit’s firing rates were z-scored and the mean z-scored firing rate for each bin was computed. For determination of significantly-modulated units (Figure 4C), similar to the minimum threshold utilized in Figure 3, we utilized a threshold of 0.75 for the difference between the maximum and minimum mean z-scored firing rates across bins.

### Data Analysis: Cell type determination

For use in the population decoding analysis described below, putative cell types were determined via similar means as previously published studies (*42, 64*) (Supplementary Figure 10). First, the template utilized by Kilosort2 to identify each particular unit was extracted and was classified as “narrow” if the width of the template was less than or equal to 400μs. Post-spike suppression indirectly measures the refractory period of a unit and was computed as the time needed for the firing rate to exceed the average firing rate in the 100-400ms period following a spike; this was computed via the auto-correlogram smoothed with a 25ms centered Hamming window. Finally, the phasic ratio was computed as the relative fraction of time a given unit spent in an interspike interval longer than 2 seconds. For units that were not considered narrow, units were deemed medium spiny neurons (MSNs) if their post-spike suppression was greater than 40ms and tonically active neurons (TANs) otherwise. For narrow units, units were deemed fast-spiking interneurons (FSIs) if their post-spike suppression was less than 40ms and their phasic ratio less than 10%. All other units were classified as unidentified interneurons and excluded from cell-type-specific analysis in Figure 4.

### Data Analysis: Population Decoding

We then utilized a population decoding model to analyze the encoding within striatal units of two particular quantities, the neural cursor and the neural velocity (Figure 4). Similar to analysis seen in Figure 1J-M, sliding windows of 10 successful trials were first computed across all sessions. Again, sessions with fewer than 10 successful trials were ignored, and only sessions within the first 12 days were considered. Firing rates spanning the 2 seconds preceding the cortical hit of each trial were extracted, and lags of −100ms and +100ms were added for each unit. Then, within each sliding window, encoding strength was assessed via cross-validated ridge regression (sklearn.linear_model.RidgeCV) fit to the trial-concatenated firing rates. In particular, 5-fold shuffled cross-validation was utilized to determine the optimal regularization constant, and all coefficients of determinations (R^2^) used in figures and models were the average test R^2^ across the 5 folds within that window. For both the neural cursor and neural velocity, a separate ridge regression model was computed within each sliding window for all striatal units and for the 6 combinations of region (dorsal and ventral striatum) and putative cell type (MSN, FSI, and TAN).

To analyze changes in encoding strength across timescales, we utilized a linear mixed model. First, as encoding strengths were highly skewed towards 1, we transformed the cross-validated R^2^ values using a log-transform to make it more amenable for linear modeling:

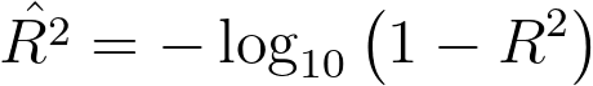

This transformation thus resulted in values between 0 and positive infinity, where more positive values represented encoding strengths closer to 1. This became our dependent variable for all population decoding models. This transformation significantly improved homoscedasticity and normality of residuals, as described below.

As done above, we first designated “early trials” as the first 15 windows of trials (i.e. spanning the first 25 trials), and “late trials” as all other windows of trials. Then, we constructed a model with fixed effects of number of days, indicator of early or late trials (−1 or 1), an interaction term between number of days and early/late indicator, and the number of units recorded that day; a random intercept per session, and a random across-day slope per rat. Critically, the inclusion of the number of units in the model accounted for inevitable increases in encoding strengths by including more predictors, and this slope was always highly significant. The models converged; while heteroscedasticity tests had p-values less than 0.05, homoscedasticity of residuals was confirmed visually for each model, and kurtosis/skew of residuals were within reasonable values (<1 for both). In some cases, the variance explained by the per-rat across-day slope was near zero, resulting in a singular random effects matrix; since this indicates there exists near-zero variation in this parameter across rats, in these cases, the per-rat across-day random slope was removed and the model re-fit.

To directly compare encoding strengths between dorsal and ventral striatum within a given cell type, we took advantage of simultaneous dorsal and ventral striatal recordings by comparing encoding strengths across all sliding windows. In particular, to account for differences in number of units across regions, we first computed the 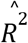 per unit recorded for each region and sliding window, and computed difference in this relative encoding strength between dorsal and ventral striatum across all sliding windows. A linear mixed model was then used to test whether this difference was significantly different than zero through a fixed effect intercept and a random intercept per session, yielding a p-value for the fixed effect intercept shown in Figure 4 (right columns for each cell type). A simpler model testing the mean difference across sliding windows per session for significant differences from zero via a 1-sample t-test yielded the same trends (data not shown).

Additionally, to analyze changes in encoding strength after the unlinking of audio and reward, we compared the encoding strengths of the five sliding windows occurring entirely and immediately before the unlinking to the five sliding windows occurring entirely and immediately after (Supplementary Figure 11). A linear mixed model with fixed effect slope and intercept and a random intercept per session was used to analyze differences in encoding strength before and after these two manipulations. All linear mixed models converged, had homoscedastic residuals, and had kurtosis/skew within reasonable bounds (< 2). Relatedly, to assess striatal encoding of the neural cursor and auditory tone frequency during catch trials, we utilized a population decoding model across the full length (~10 seconds) of all catch trials within a given session, and compared encoding strengths with a paired t-test.

To analyze alternative hypotheses of striatal encodings, we compared the encoding strength of the neural cursor with that of a linear representation of time, as striatum has been reported to be correlated with measures of time (*65*) (Supplementary Figure 12). We considered the same time periods as previous population decoding analyses (2 seconds before hit), except models were fit to an entire session’s trials, instead of windows of trials, to simplify analysis. As a measure of time, a line was constructed that ramped from 0 to the max value of the mean neural cursor, starting at an onset swept across a range of values (−2s, −1.5s, −1.4s, −1.0s, −0.6s, −0.5s, −0.4s, −0.3s, and −0.2s relative to hit). Then, in order to enable a fair comparison of encodings strengths with the neural cursor, each representation of time was variance-matched to the neural cursor: per-time-bin variance was computed for the neural cursor across all successful trials, and Gaussian noise with equal variance was injected into the line. A population decoding model was then fit using units within the respective subgroups of striatal region and cell types. We then only retained the onset for the time representation that had the highest encoding strength. Lastly, similar to Supplementary Figure 11, a linear mixed model with fixed effect slope and intercept and a random intercept per session was used to assess differences in encoding strengths.

Similarly, as an alternative hypothesis to neural velocity encodings, striatum could have encoded a discrete “go” signal beginning at some point before hit, corresponding to a step function (Supplementary Figure 12). We performed a similar analysis as investigating time, except a step function was used, instead of a line, and amplitudes/variances were matched to the neural velocity signal instead of neural cursor.

## Supplementary Figures and Data

**Supplementary Figure 1.**
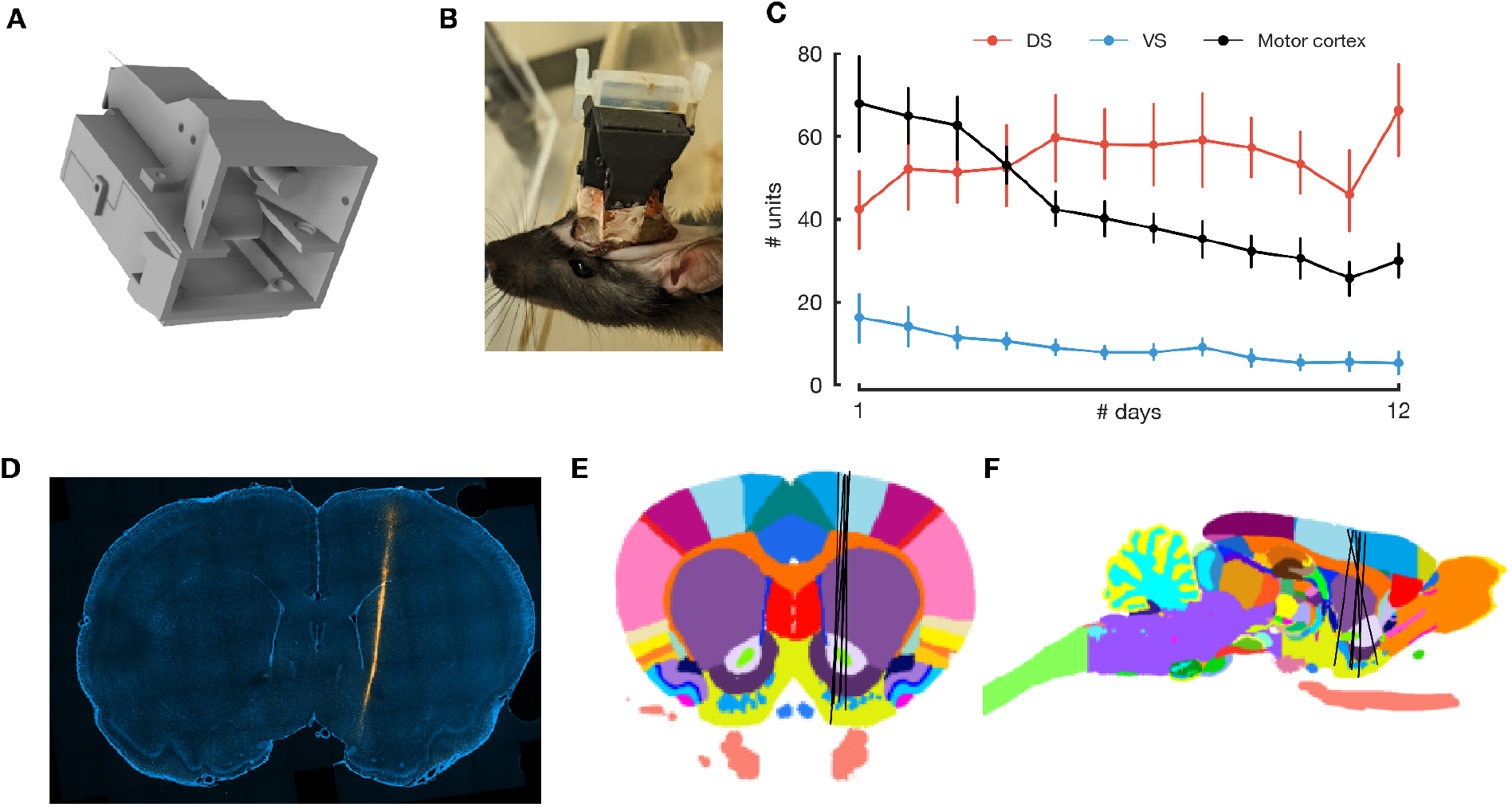
Custom-made implant yields tens to hundreds of units each day, and histology confirms placement of probe within targeted areas of striatum. **A:** 3D CAD model of assembled custom-made 3D-printed implant, heavily based on a prior validated design (*40*). View is from the end of the implant that will be facing dorsally. The Neuropixels 1.0 headstage PCB is secured via M1 screws in the bottom half of the implant (as viewed from this angle); the cylinder visible in the top half is used to attach the implant to the stereotactic frame via a cannula holder. **B:** entire implanted probe assembly on a ~250g rat after a few days of recovery post-surgery. Black and transparent white material is the 3D-printed implant; orange wrapping is copper foil used for shielding. **C:** number of well-isolated units recorded per day, divided into regions as defined in Methods; bars represent SEM. **D:** example stained and imaged histological brain slice. Slice was stained with DAPI (blue); probe track (orange) fluoresces from Vybrant DiI dye applied to the probe shank before implantation. **E, F:** reconstructed probe tracks (n=6) overlaid on coronal (**E**) and sagittal (**F**) planes of the Waxholm Space Atlas. In 1 rat VS units were excluded due to posterior location of probe (see methods). Different colors represent the default colors used to differentiate atlas-defined brain regions.

**Supplementary Figure 2.**
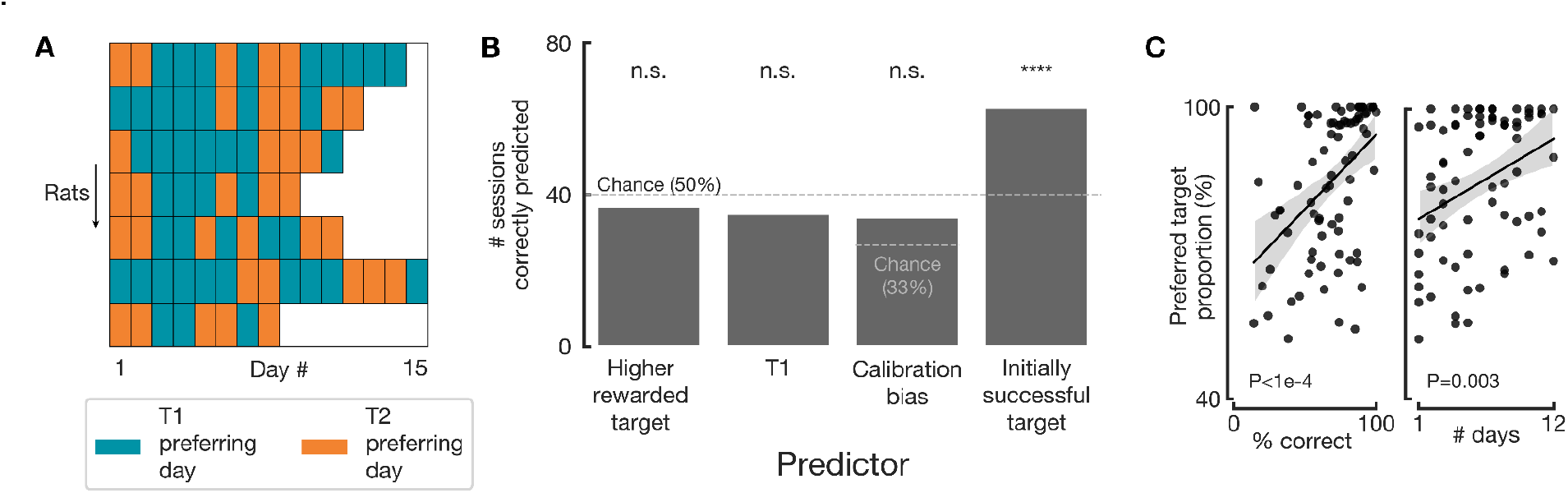
Rats preferred the target they initially achieved. **A:** display of all included sessions (n=80) and which target (T1 or T2) was more highly preferred that day, i.e. the target that had more successful trials. Each row represents a single rat, and each box a day. In total, 86.8% of trials went to the preferred target for that day. **B:** measures of possible factors contributing to a rat choosing a particular target on a particular day. Rats did not prefer the target leading to more reward (binomial test for 37/80, p=0.58), did not consistently prefer a particular target overall (T1 chosen arbitrarily; binomial test for 35/80, p=0.31), and did not prefer the target that might have been biased due to calibration (bias determined as the target with more rewards in simulated calibration; each (neuroprosthetic) session was then determined as not biased or biased according to a binomial test p-value of less than 0.05; final predictor significance determined via binomial test for 34/80 with 3 choices, p=0.10). However, rats did significantly prefer whichever target they achieved more frequently within the first 5 trials (binomial test for 63/80, p<1e-6). **C:** The proportion of trials per session going to a preferred target correlates significantly with the proportion of correct trials (left, p=1.9e-5) and with time (right, days #1-12 only, p=0.003). Individual dots represent 1 session; black line and gray shading indicate regression line and lowess, respectively.

**Supplementary Figure 3.**
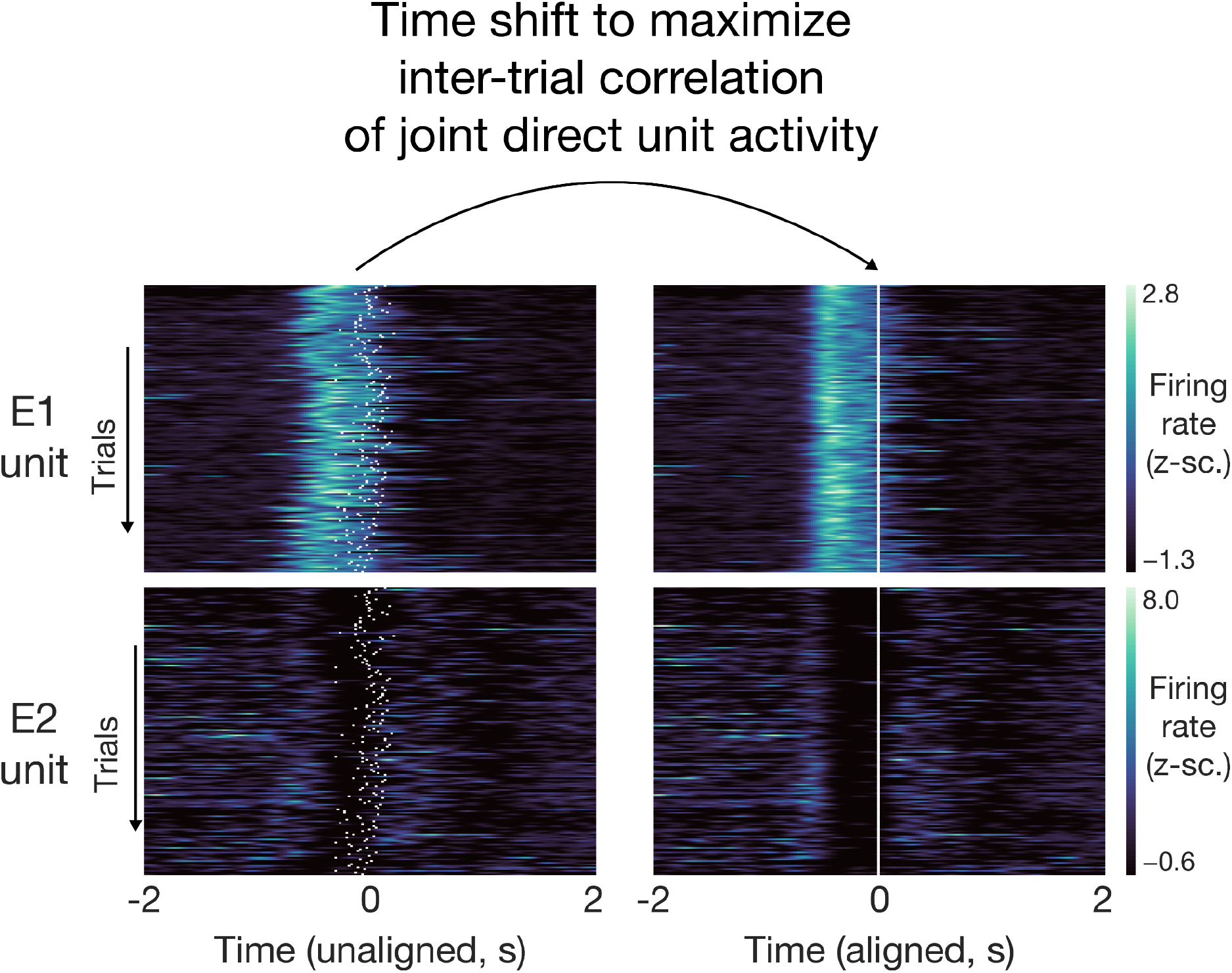
Demonstration of time-shifting utilized to align trials, correcting for trial-by-trial variations in precise timing. Optimal time shifts were computed via previously demonstrated algorithms (*57*) that optimized the inter-trial correlations of direct unit activity for successful trials of each target. Left: unaligned, simultaneous firing rates for a unit in ensemble 1 (top) and a unit in ensemble 2 (bottom). White ticks indicate the model’s prediction of the alignment point and will be the points that become “time = zero” after alignment. Right: aligned, simultaneous firing rates for the same units and trials. All other units’ activities (not shown) were then shifted by the same amount as the direct units to maintain alignment.

**Supplementary Figure 4.**
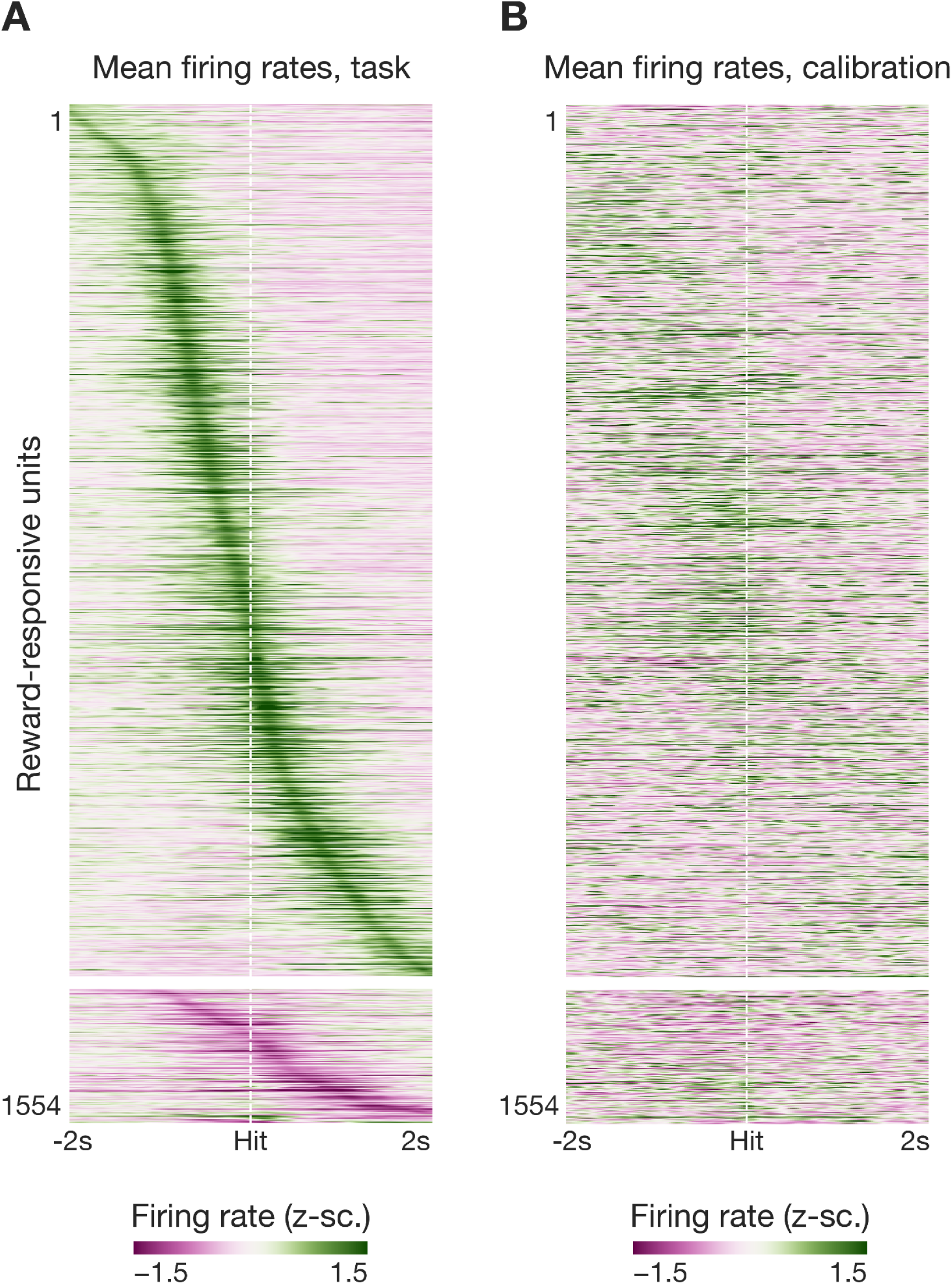
Mean firing rates for reward-responsive units time-locked to target hit during the task (**A**) and to simulated target hit during each task’s preceding calibration (**B**). Units are the same across rows. Firing rates were z-scored independently between task and calibration.

**Supplementary Figure 5.**
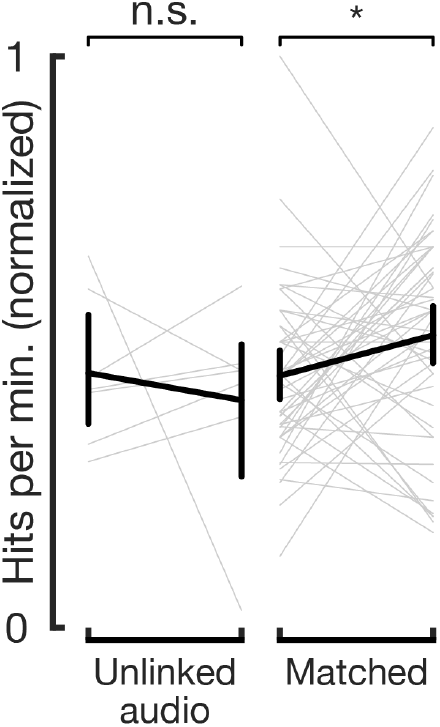
Performance does not change when the auditory tone is unlinked from cortical activity. In a subset of high-performing, late-learning sessions, the frequency of the auditory tone was unlinked from cortical activity and instead linearly ramped across the frequency spectrum, while the link between cortical activity and reward remained intact. There was no significant change in the hits per minute between the 10 minutes preceding and the 10 minutes following unlinking of audio (n=7 sessions, p=0.66). In performance-matched, normal sessions, there was a slight but significant increase in performance in a similar time period (n=48, p=0.04; same as pictured in Figure 1I).

**Supplementary Figure 6.**
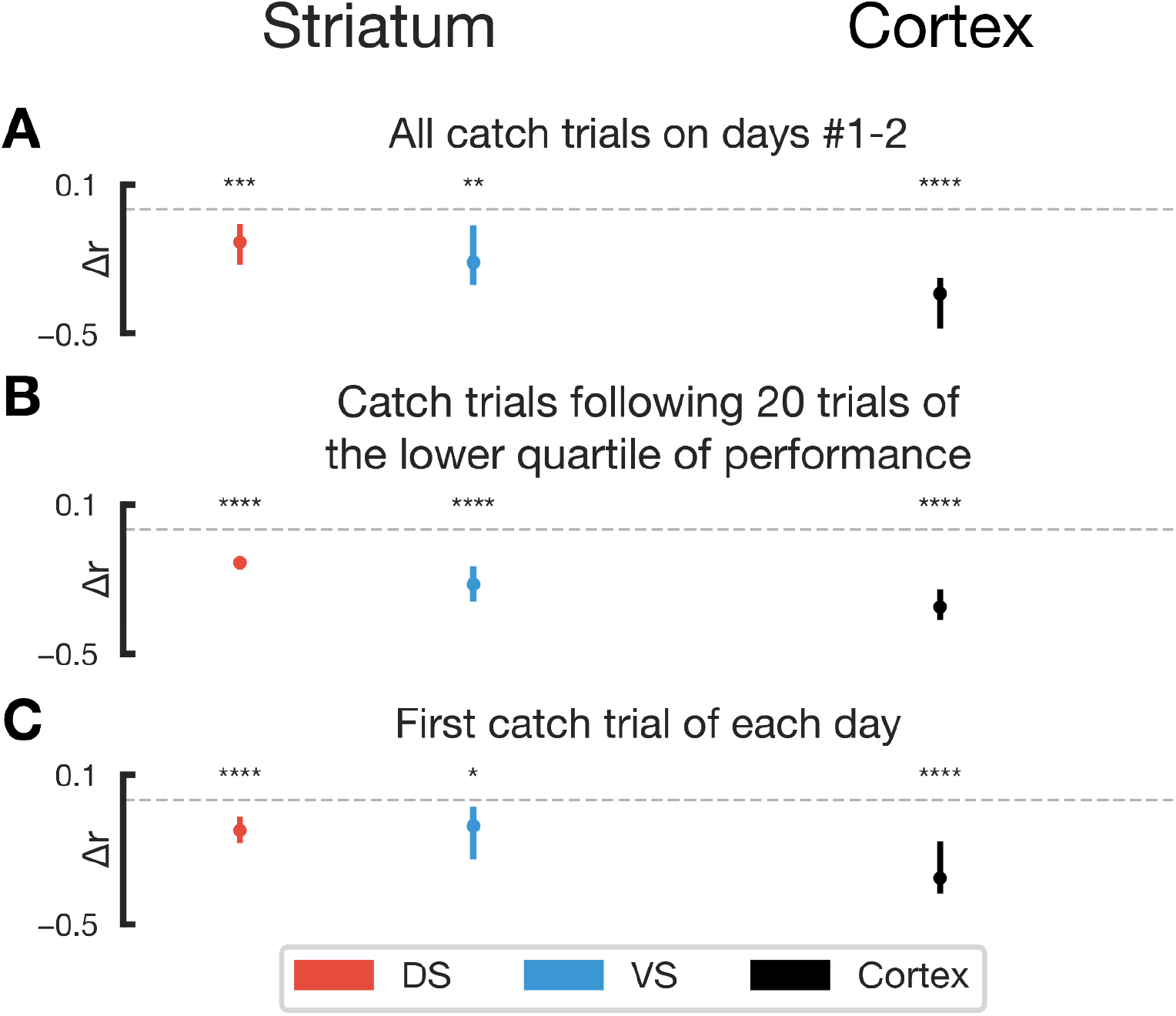
Further analysis of striatal representations at the end of catch trials. Display and analysis is the same as that of Figure 3, with analysis performed on different subsets of catch trials. **A:** trial-to-trial template correlations computed at the end of all catch trials only on the first 2 days. **B:** correlations analysis performed on catch trials following a period of 20 trials that were in the lowest 25th percentile of mean hit times, representing catch trials following poor performance. **C:** correlations analysis for the first catch trials of each day. For all, n.s. p > 0.05; * p < 0.05, ** p < 0.01, *** p < 0.001, **** p < 0.0001.

**Supplementary Figure 7.**
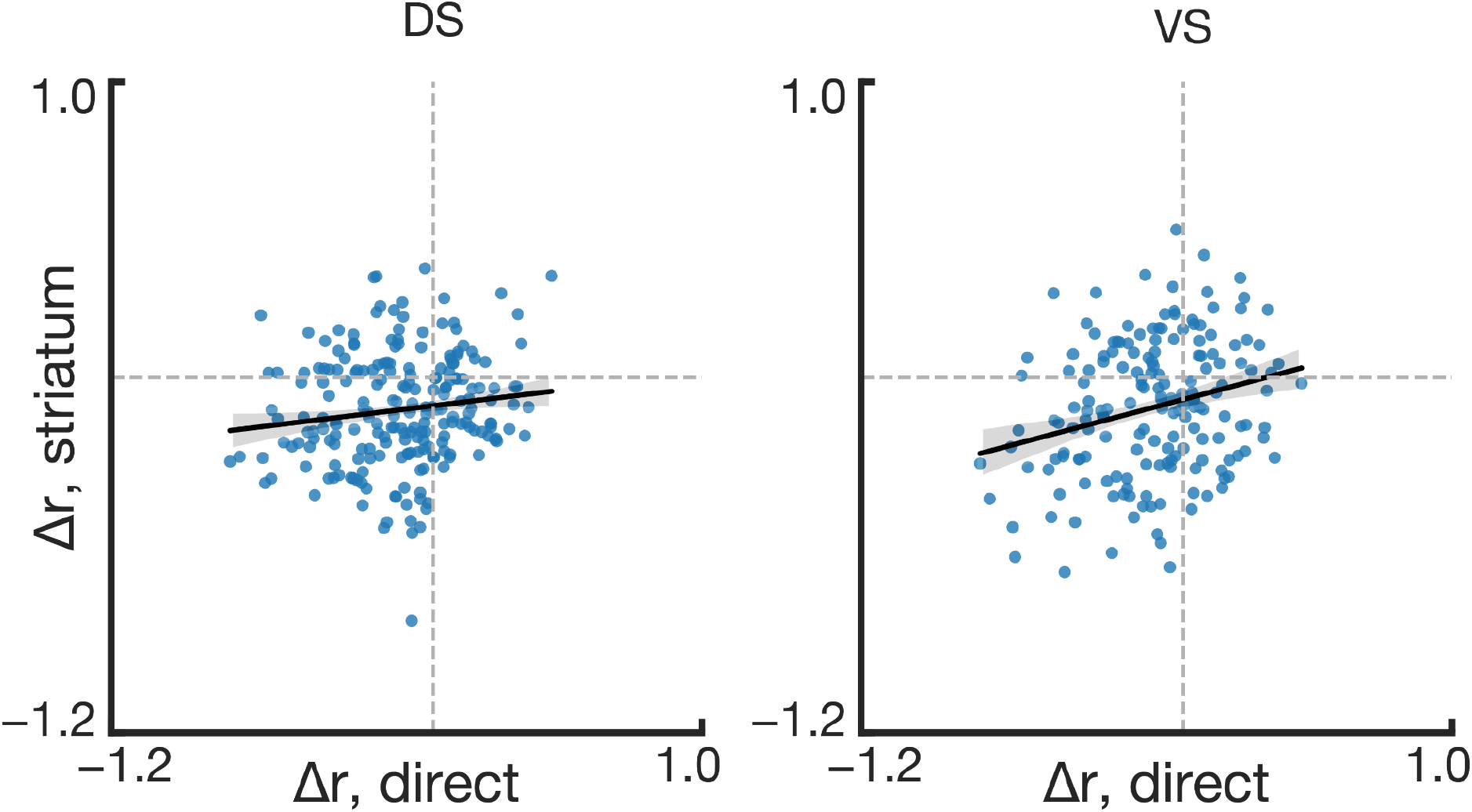
Further analysis of striatal representations during outside-trial hits. For each outside-trial hit analyzed, the median Δr value for all dorsal (left) / ventral (right) striatal units (x-axis) was plotted against the Δr value for the direct units (y-axis), and a linear regression model was fit (black line: regression line, shading lowess). The intercepts for both dorsal and ventral striatum were significantly negative (DS −0.09, p<0.0001; VS −0.07, p<0.001), indicating that even when cortical activity in extra-trial hits is similar to preceding successful trials, striatal activity still significantly deviates from its current representation.

**Supplementary Figure 8.**
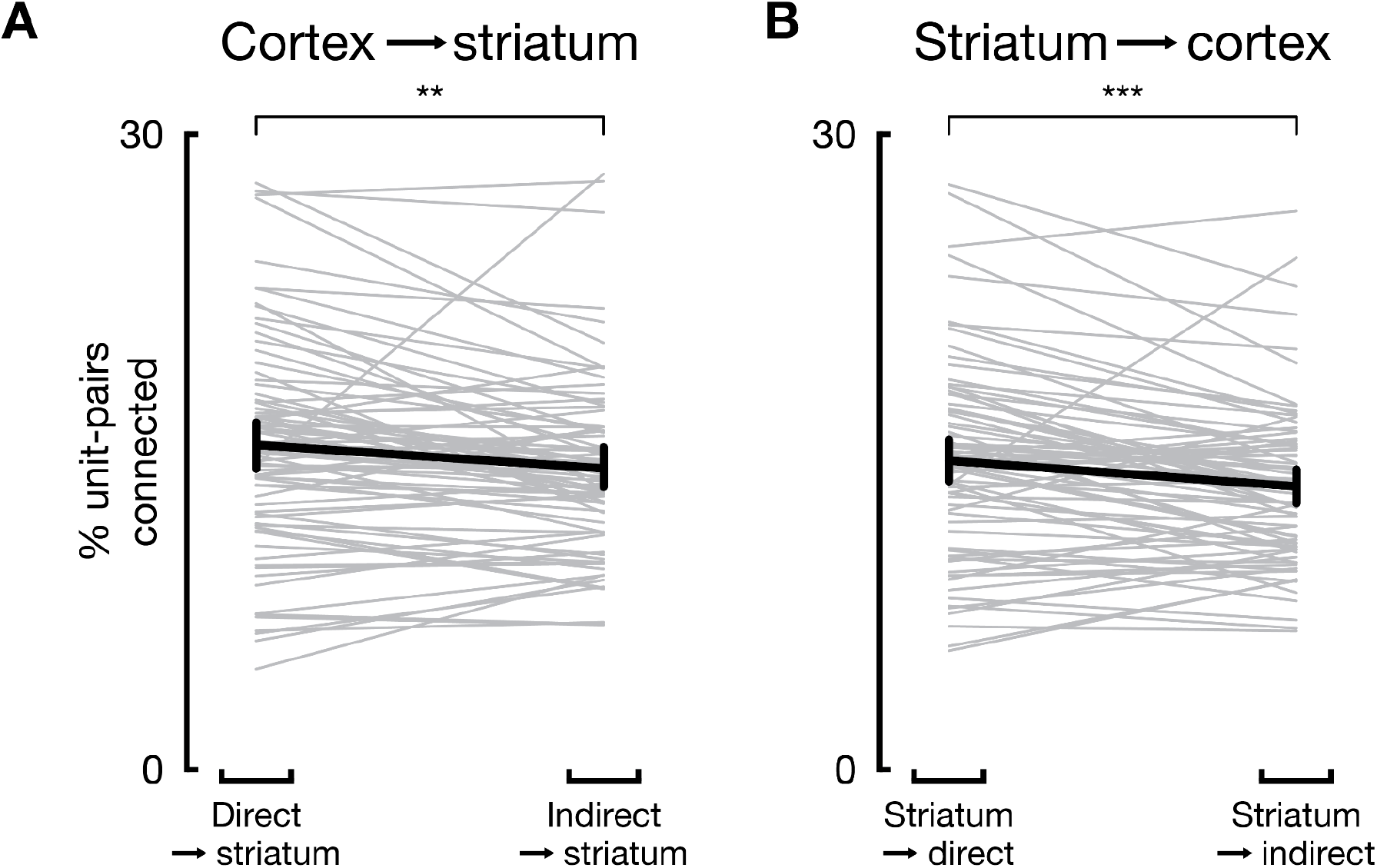
Granger causality analysis between striatal and motor cortical units shows functional connectivity is higher between striatum and direct units than striatum and indirect units. **(A)** Percent of unit-pairs connected with direct (left) or indirect (right) cortical units as the source, and striatal units as the targets (p=0.002, paired t-test). **(B)** Same as A but with source and target reversed (p = 0.0003, paired t-test). For all, * p < 0.05, ** p < 0.01, *** p < 0.001, **** p < 0.0001, otherwise not indicated p > 0.05; bars represent 95% confidence intervals.

**Supplementary Figure 9.**
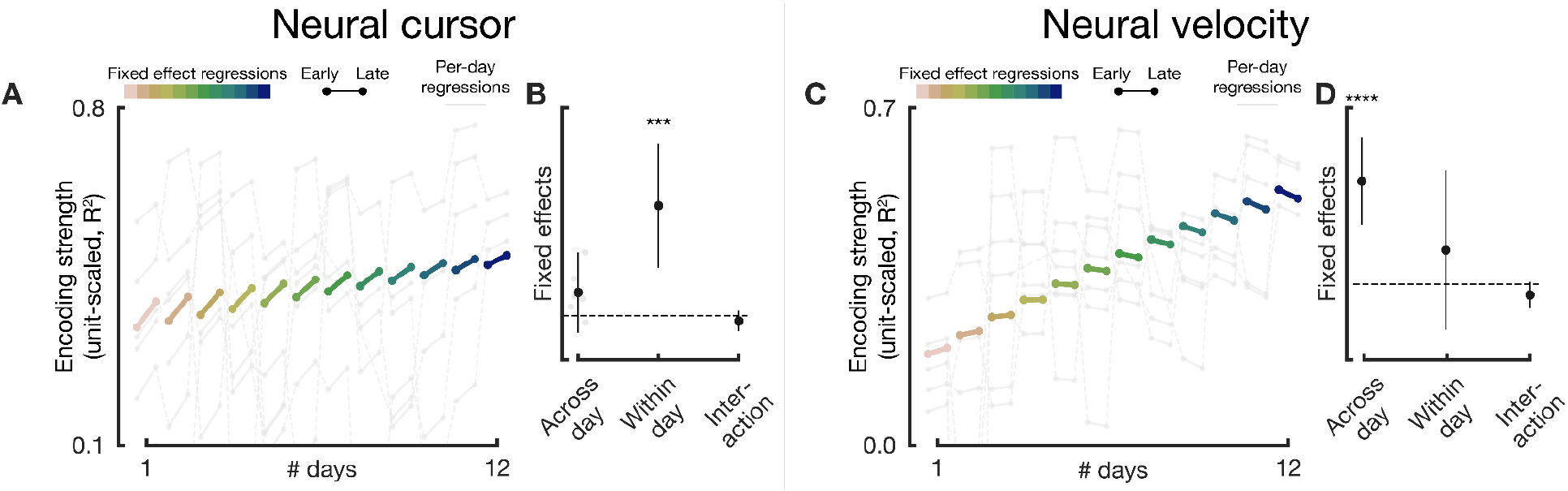
Results of population decoding model when all striatal units are included for neural cursor encoding (left) and neural velocity encoding (right). **(A)** Regressions from linear mixed model (fixed effects: colored lines; per-day regressions incorporating random effects: gray lines) displaying changes in neural cursor encoding strength both within-day (early vs late trial windows, indicated by endpoints of each line) and across-day changes (x-axis). Encoding strengths shown are scaled to 20 units (“unit-scaled”, Methods). Model intercept for unit-scaled R^2^ 0.37 (p<1e-4). **(B)** Fixed effect slopes for neural cursor encoding; positive slopes indicate increases in encoding strength. Faded gray dots indicate random effects, when applicable. Y-axis for fixed/random effects is log-transformed R^2^ (Methods). **(C-D)** Same as (A) and (B), except with neural velocity encoding. Model intercept for unit-scaled R^2^ 0.21 (p<1e-6). For all, * p < 0.05, ** p < 0.01, *** p < 0.001, **** p < 0.0001, otherwise not indicated p > 0.05; bars represent 95% confidence intervals.

**Supplementary Figure 10.**
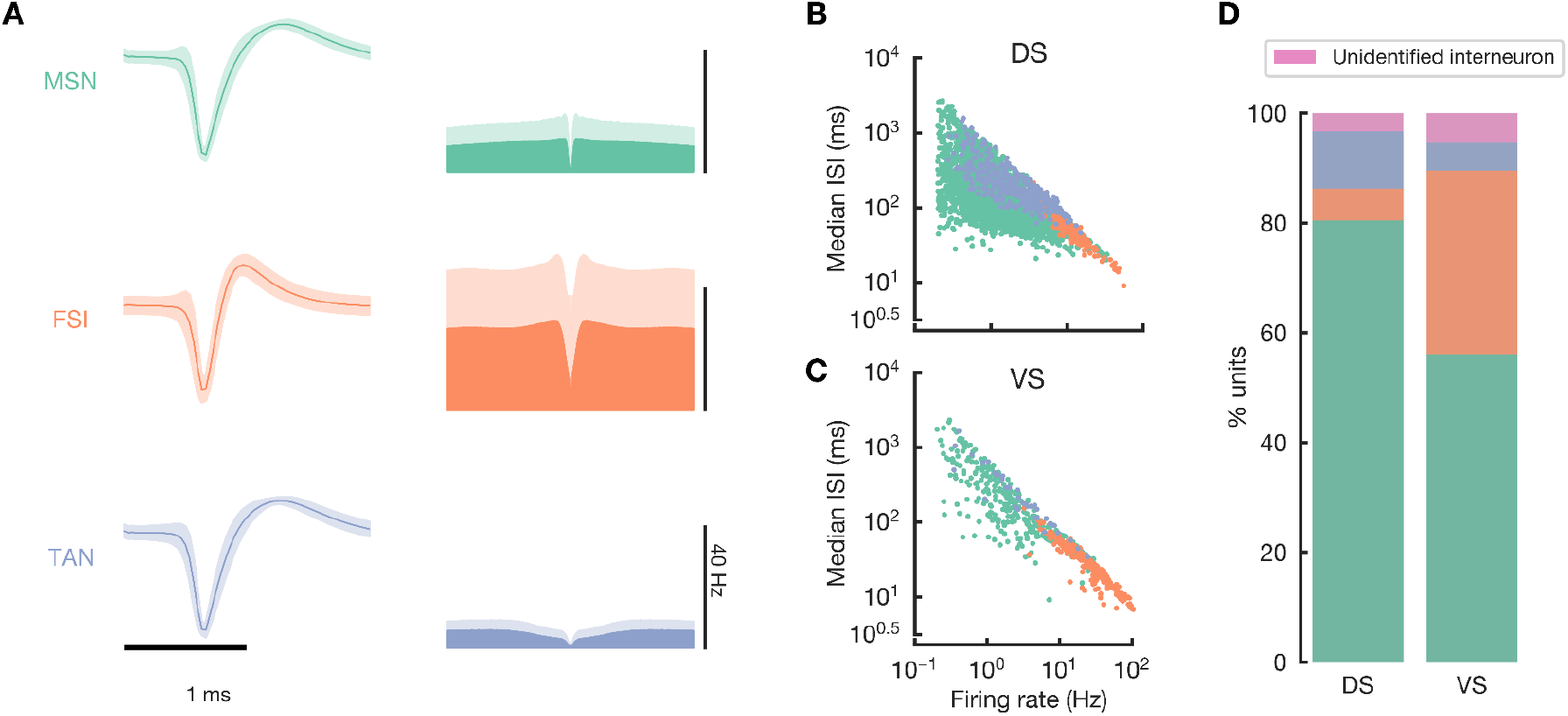
Classification of units into putative cell types via electrophysiological characteristics. **A:** left: mean template extracted by Kilosort2 for all units classified as a particular cell type (MSN: medium spiny neuron, FSI: fast-spiking interneuron, TAN: tonically active neuron). Shading is 1 standard deviation. Right: mean auto-correlograms for each cell type, displayed from −150ms to 150ms (x-axis). Shading is 1 standard deviation. **B:** distribution of firing rate properties for all well-isolated units within the dorsal striatum identified as particular cell types. Colors are the same as in **(A)**; ISI interspike interval. **C:** same as (B) but for units in ventral striatum (VS). **D:** relative proportion of each cell type within the population of well-isolated dorsal and ventral striatal units. 3943 dorsal striatal units were identified as 3174 MSNs, 228 FSIs, 411 TANs, and 130 unidentified interneurons; 690 ventral striatal units were identified as 387 MSNs, 231 FSIs, 35 TANs, and 37 unidentified interneurons.

**Supplementary Figure 11:**
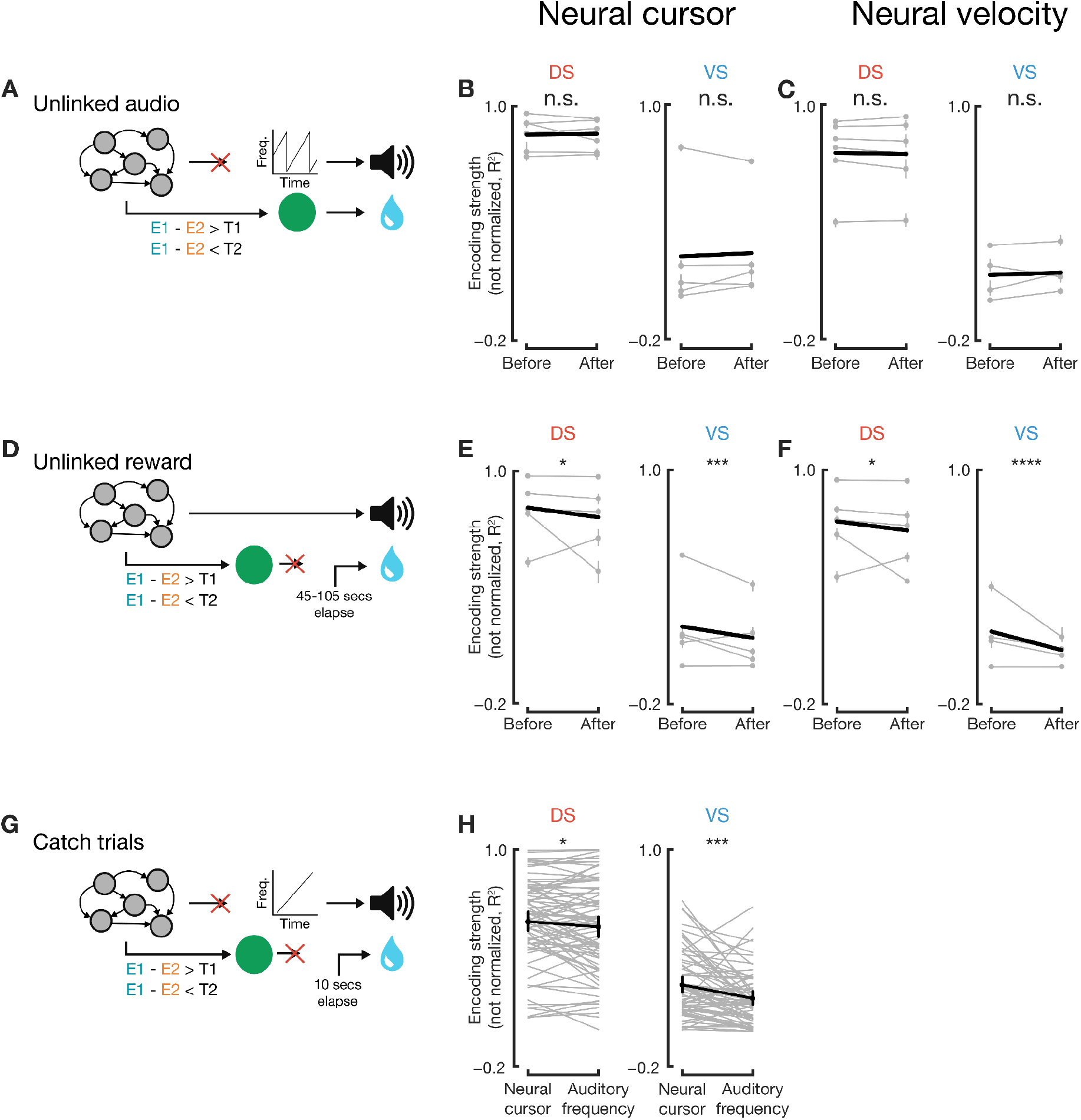
Population decoding model applied to instances where audio and reward were unlinked and during catch trials, as also depicted at the single-unit level in Figure 3.. **A:** when the auditory tone was unlinked from cortical activity in a subset of sessions, there were no significant changes in the encoding strength within dorsal or ventral striatum for the neural cursor or velocity (p=0.67, p=0.20, p=0.55, p=0.39 for dorsal-cursor (B, left), ventral-cursor (B, right), dorsal-velocity (C, left), ventral-velocity (C, right) encodings, respectively). **D:** when reward was unlinked from cortical activity, encoding strength for both neural cursor and velocity significantly decreased in both dorsal and ventral striatum (p=0.03, p=0.0003, p=0.02, p=0.0001 for dorsal-cursor (E, left), ventral-cursor (E, right), dorsal-velocity (F, left), ventral-velocity encodings (F, right), respectively). **G:** during catch trials, the auditory tone and the neural cursor were decoupled.Both dorsal (H, left) and ventral striatum (H, right) encoded the neural cursor more strongly than the auditory tone’s frequency (DS p=0.03, n=80 sessions, VS=0.0005, n=67 sessions; paired t-test). In all, thin gray lines represent individual session’s data; thick black lines indicate regressions. Bars indicate 95% confidence interval.

**Supplementary Figure 12:**
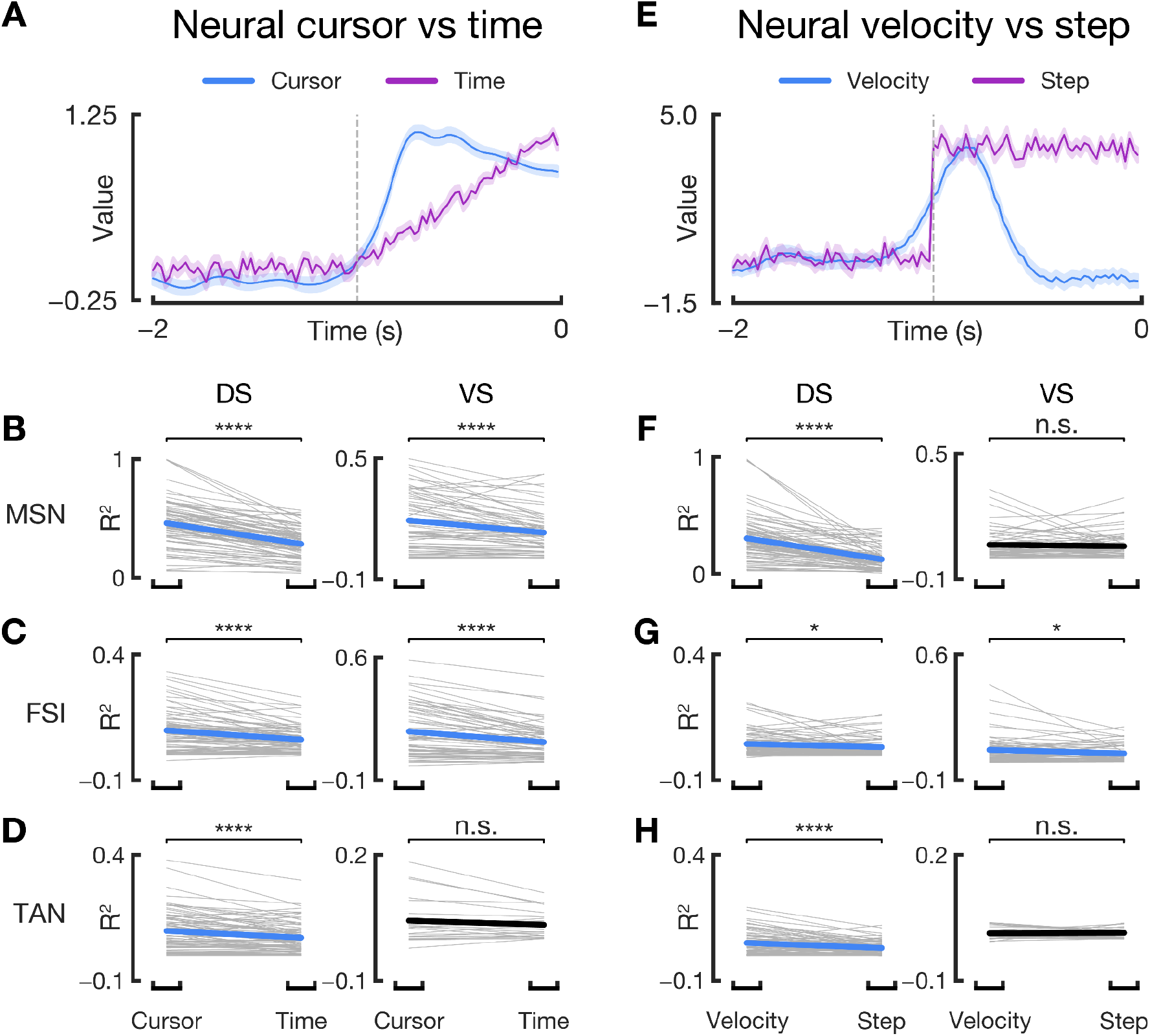
Population decoding model applied to alternative encoding hypotheses, where striatal activity may encode a linear representation of time (*62*) instead of the neural cursor (left column), or may encode the time derivative of this representation of time, a step function, instead of the neural velocity (right column). **(A)** Example neural cursor and time data. Time (purple) functions were constructed based on a linear ramp from 0 to the max value of the mean neural cursor, where the onset time (gray dotted line) was swept across a range of times. Additionally, Gaussian noise was injected into the time function to match the variance of the neural cursor to enable a fair comparison for encoding. Shading SEM. **(B-D)** Comparison of encoding strengths (R^2^, not unit-scaled) across an entire session for the encoding of the neural cursor (left datapoint of each column) and time (right datapoint of each column) for all 6 combinations of regions & putative cell types. For each session, the best onset for the time function was taken. Significance of the slope (i.e. the difference between cursor and time encoding strengths) was computed via a linear mixed model. Bold line indicates fixed effect; black bold line means slope is not significant, otherwise the color represents which quantity is more strongly encoded. Gray lines represent each session. As in other figures: n.s. p > 0.05; * p < 0.05, ** p < 0.01, *** p < 0.001, **** p < 0.0001. **(E)** Similar to (A), a step function was constructed, with scale equal to the max mean neural velocity value and onset time swept across a range of times. Variances were matched as well. Data originates from the same session as (A). **(F-H)** Same as (B-D), but comparing neural velocity (left datapoint) and step function (right datapoint) encodings.

